# MuSK-BMP signaling in adult muscle stem cells maintains quiescence and regulates myofiber size

**DOI:** 10.1101/2023.05.17.541238

**Authors:** Laura A. Madigan, Diego Jaime, Isabella Chen, Justin R. Fallon

## Abstract

A central question in adult stem cell biology is elucidating the signaling pathways regulating their dynamics and function in diverse physiological and age-related contexts. Muscle stem cells in adults (Satellite Cells; SCs) are generally quiescent but can activate and contribute to muscle repair and growth. Here we tested the role of the MuSK-BMP pathway in regulating adult SC quiescence by deletion of the BMP-binding MuSK Ig3 domain (‘ΔIg3-MuSK’). At 3 months of age SC and myonuclei numbers and myofiber size were comparable to WT. However, at 5 months of age SC density was decreased while myofiber size, myonuclear number and grip strength were increased - indicating that SCs had activated and productively fused into the myofibers over this interval. Transcriptomic analysis showed that SCs from uninjured ΔIg3-MuSK mice exhibit signatures of activation. Regeneration experiments showed that ΔIg3-MuSK SCs maintain full stem cell function. Expression of ΔIg3-MuSK in adult SCs was sufficient to break quiescence and increase myofiber size. We conclude that the MuSK-BMP pathway regulates SC quiescence and myofiber size in a cell autonomous, age-dependent manner. Targeting MuSK-BMP signaling in muscle stem cells thus emerges a therapeutic strategy for promoting muscle growth and function in the settings of injury, disease, and aging.

**Highlights:**

- MuSK, in its role as a BMP co-receptor, regulates adult muscle stem cell quiescence
- The MuSK-BMP pathway acts cell autonomously
- Increased muscle size and function with preservation of myonuclear density and stemness in mice with attenuated MuSK-BMP signaling

## INTRODUCTION

Adult muscle stem cells (Satellite Cells; SCs), are key arbiters of muscle growth, homeostasis and regeneration(Keefe et al., 2015; Murach et al., 2021; Pawlikowski et al., 2017; Sambasivan et al., 2011; Sousa-Victor et al., 2022). SC function is tailored to the state of the muscle as well as to age. For example, during the first weeks of life large scale SC proliferation and fusion are essential for achieving mature muscle size and myonuclear complement. In mice, SCs contribute to normal myofiber growth until ∼8 weeks of age(Bachman et al., 2018; Bachman and Chakkalakal, 2022; Cramer et al., 2020; Gattazzo et al., 2020; Pawlikowski et al., 2015; White et al., 2010). After this age SCs in uninjured muscles are generally quiescent, but can activate to support muscle growth in response to load or exercise(Egner et al., 2016; Goh et al., 2019; Goh and Millay, 2017; Murach et al., 2021). SCs can also activate and proliferate following injury to effect muscle regeneration(Murphy et al., 2011; Sambasivan et al., 2011; Zammit et al., 2002). Finally, dysregulation of SC function with aging is thought to contribute to sarcopenia(Blau et al., 2015; Brack et al., 2005; Sousa-Victor et al., 2022).

A key question in SC biology is elucidating the signaling pathways that regulate their dynamics, particularly in age-related contexts. Known pathways include Notch-Delta, Wnts, Collagens V and VI, Cadherin and BMPs(de Morree and Rando, 2023; Fuchs and Blau, 2020; Kann et al., 2021). Notch signaling controls both satellite cell quiescence and is necessary for SC self-renewal following activation(Bjornson et al., 2012; Gioftsidi et al., 2022; Zhang et al., 2021). Muscle-derived Wnt4 maintains SC quiescence(Eliazer et al., 2019). Members of the TGFß superfamily also play important roles. Myostatin (GDF8) is a potent regulator of SC dynamics during early postnatal development, with its absence resulting in massive increases in muscle size. However, myostatin does not regulate SC activation in adults(Lee, 2022; Murphy et al., 2010). TGFß signaling is a negative regulator of myoblast fusion(Girardi et al., 2021). Bone morphogenetic proteins (BMPs), regulate cell proliferation and subsequent SC-mediated muscle growth in both development and during regeneration(Gozo et al., 2013; Ono et al., 2011; Stantzou et al., 2017). However, a role for BMP signaling in SC quiescence in adult muscle has not been described.

We recently reported that the transmembrane protein MuSK, in addition to its well-established role as an agrin-LRP4 receptor, is also a BMP co-receptor that shapes BMP-mediated transcriptional output in myogenic cells(Fish and Fallon, 2020; Jaime et al., 2024; Yilmaz et al., 2016). MuSK binds to BMPs as well as to the type I BMP receptors BMPR1a and BMPR1b (also termed ALK3 and ALK6, respectively). Cultured myogenic cells expressing MuSK show enhanced pSmad1/5/8 signaling and express a distinct set of transcripts in response to BMP stimulation compared to MuSK^-/-^ cells. MuSK-dependent BMP signaling is dependent on Type I BMP receptor activity, but MuSK tyrosine kinase activity is dispensable. Importantly, the Ig3 domain of MuSK is necessary for high affinity BMP binding, while this region is not required for agrin-LRP4 signaling (which is mediated by the Ig1 domain). The role of MuSK as a BMP co-receptor is thus structurally and functionally distinct from its function in agrin-LRP4 signaling. We recently created a germline knock-in mouse model lacking the MuSK Ig3 domain(Jaime et al., 2024). These ‘ΔIg3-MuSK’ mice are viable and fertile and survive to at least 27 months of age. As predicted, myogenic cells isolated from these mice show reduced pSMAD and transcriptional responses to BMP, but normal agrin-LRP4-dependent activity. In that study we characterized these mice at 3 months of age and showed that the MuSK-BMP pathway maintains muscle size in the slow soleus muscle (but not in the fast TA) by regulating myofiber-intrinsic Akt-mTOR signaling.

Here we investigated the role of the MuSK-BMP pathway in SCs in both uninjured and regenerating fast TA and EDL muscles. We show that MuSK transcript and protein are expressed in WT and ΔIg3-MuSK SCs in uninjured muscle. At 3 months of age the ΔIg3-MuSK TA shows normal myofiber size and SC numbers. However, at 5 months SCs are activated and myonuclear number, myofiber size, muscle weight and grip strength are elevated. Notably, 5- month-old mutant muscles efficiently regenerate after injury and the SC pool and myofiber size is restored to WT levels, indicating that the ΔIg3-MuSK SCs maintain full stem cell function. Finally, conditional mutants established that the role of MuSK in regulating quiescence is SC-autonomous and is active in the adult. We conclude that MuSK expressed in SCs regulates quiescence and muscle growth in an age-dependent manner via its role as a BMP co-receptor.

## RESULTS

### MuSK is expressed in satellite cells

Previous transcriptomic analyses have detected MuSK mRNA in quiescent, activated and regenerating SCs(Petrany et al., 2020; Shcherbina et al., 2020; Yue et al., 2020). We confirmed that MuSK transcripts are expressed in SCs isolated from uninjured and regenerating muscle from both 3- and 5- month old mice using gel-based PCR analysis (**Fig. 1a-c; Supp. Fig 1a, b**). This analysis also showed that all MuSK detected in WT SCs harbors the Ig3 domain (encoded by exons 6 and 7). For clarity we refer to the form containing the Ig3 domain as Full-Length (FL)-MuSK. We did not detect transcripts lacking the Ig3 domain (ΔIg3-MuSK) in isolated WT SCs (**Fig. 1c**). To detect MuSK protein in SCs we isolated myofibers from adult muscle and double labeled them with antibodies recognizing Pax7 and MuSK. As shown in **Fig. 1d**, MuSK protein is expressed in SCs from both WT and ΔIg3-MuSK mice.

**Figure 1.**
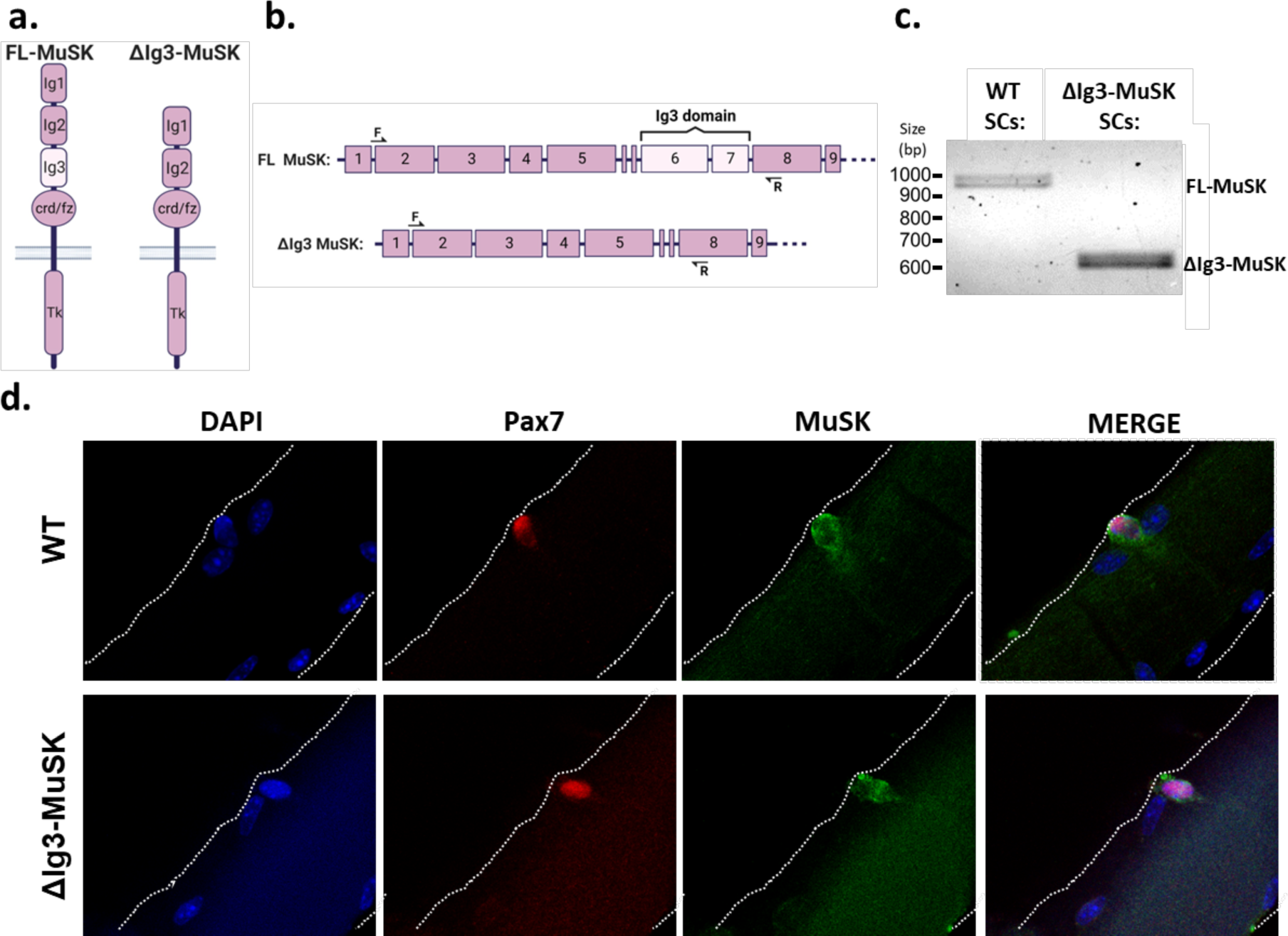
MuSK is expressed in SCs from WT and ΔIg3-MuSK mice. (**a**) MuSK protein domain schematic. Full-length (FL)-MuSK contains the Ig3 domain, while it is lacking in ΔIg3-MuSK. (**b**) Schematic of MuSK exon organization including approximate locations of forward (F) and reverse (R) PCR primers used for detection of either FL-MuSK, containing exons 6 & 7 (which encode the Ig3 domain) or ΔIg3-MuSK, lacking these exons. Note that the presence of two small (30nt) alternatively-spliced exons, ‘5a’ and ‘5b’ result in multiple bands (See also Supplemental Fig. 1). **(c)** PCR analysis of FACs-sorted SCs: WT SCs express FL-MuSK; endogenous ΔIg3-MuSK is not detected. SCs from ΔIg3-MuSK mice express only the form lacking exons 6 and 7. **(d)** IHC of isolated WT and ΔIg3-MuSK adult EDL myofibers cultured for 1d and stained for nuclei (DAPI), SCs (Pax7) and MuSK (see Methods). Note that MuSK is expressed in SCs of both genotypes. White dotted line denotes myofiber perimeter.

### Myofiber diameter and SC numbers are comparable in 3-month-old WT and ΔIg3-MuSK TA muscle

As a first step towards assessing the role of the MuSK-BMP pathway in SC dynamics we compared both SC number and myofiber diameter in WT and ΔIg3-MuSK TA from 3-month-old mice. We chose this time point since adult SC number and myonuclear content are established by this age and SCs are in their adult quiescent state(Ancel et al., 2021; Bachman and Chakkalakal, 2022; Cramer et al., 2020). Analysis of Pax7+ cells showed that their density was comparable in both genotypes at 3 months (**Fig. 2a, c**). TA myofiber diameter was also comparable in both genotypes at this age (**Fig. 2a, b**), which is in agreement with results in a related study(Jaime et al., 2024). These results indicate that the MuSK-BMP pathway does not play a discernable role in the developmental and early postnatal events that establish myofiber size and SC pool numbers in fast muscle.

**Figure 2.**
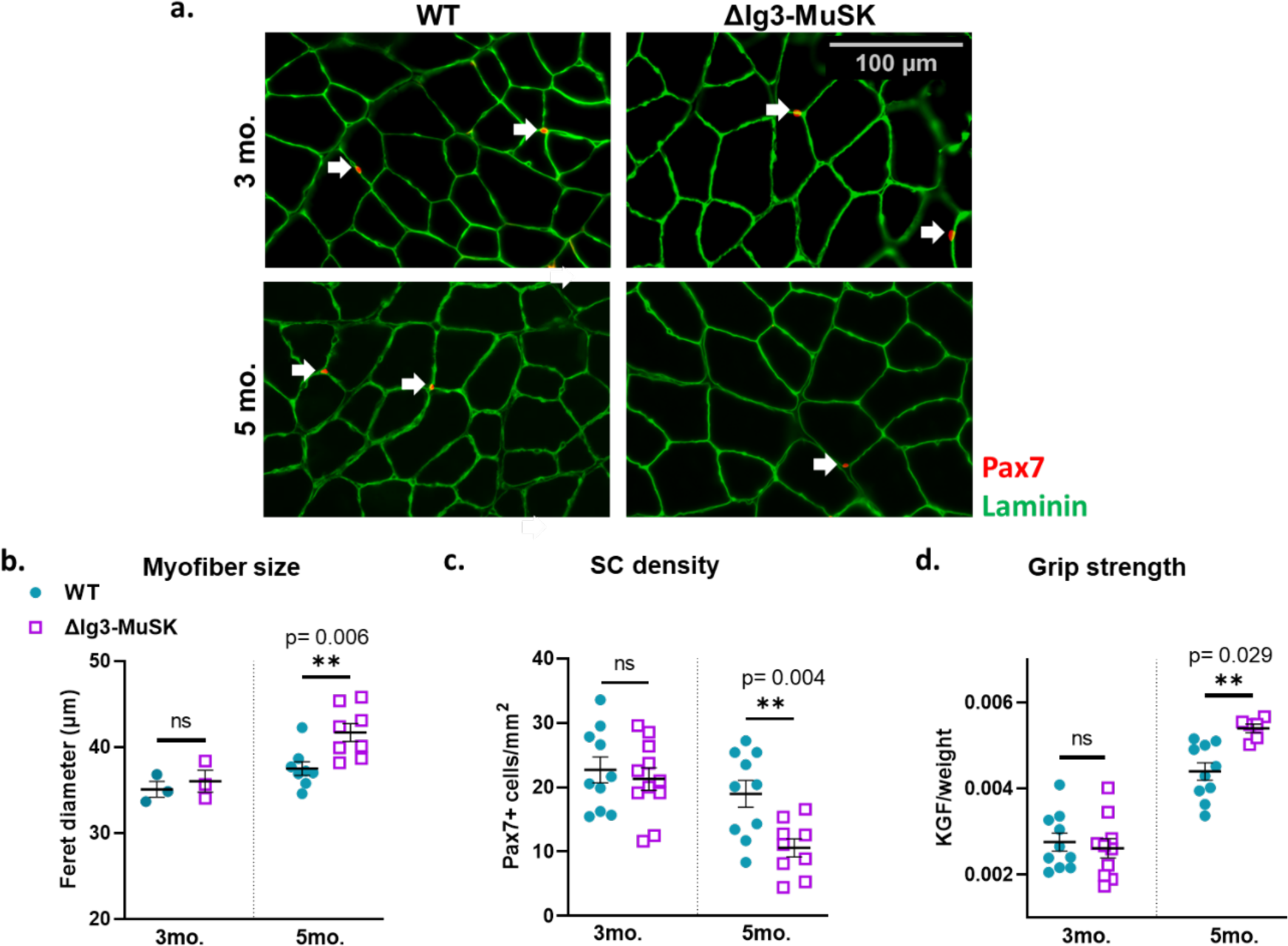
Changes in TA myofiber size, SC number, and grip strength emerge between 3 and 5 months of age in ΔIg3-MuSK mice. (**a**) Representative IHC images of 3- and 5-month-old WT and ΔIg3-MuSK TA used for Pax7+ quantification and myofiber size analysis. Pax7 (red) and laminin (green) staining; arrows indicate SCs. (**b**) At 3 months, myofiber minimum Feret diameter is equivalent in WT and ΔIg3-MuSK muscle (n=3 male mice, unpaired t-test, 3 mo. p=0.6). At 5 months, myofiber minimum Feret diameter is increased by about 10% (37 ±0.80µm and 41 ± 1.0µm in WT vs ΔIg3-MuSK, respectively; n=8 male mice, unpaired t-test p=0.006). (**c**) At 3 months of age, the number of Pax7+ cells per area is equivalent between WT and ΔIg3-MuSK muscle (n=10-11 male & female mice, unpaired t-test, p=0.6). At 5 months, the number of Pax7^+^ cells per area is reduced by almost half in ΔIg3-MuSK muscle (19±2.1 and 10.6±1.4mm^2^ in WT vs ΔIg3-MuSK, respectively; n=9-10 male & female mice, unpaired t-test, p=0.004). (**d**) At 3 months, grip strength is comparable between the two genotypes (n=10 males, unpaired t-test, p=0.64). In contrast, at 5 months, ΔIg3-MuSK mice have increased grip strength (n=6-10 males, unpaired t-test, p=0.029; see also Supplemental Fig. 2).

### Myofiber diameter is increased while SC numbers are reduced in 5-month-old ΔIg3-MuSK TA muscle

To determine whether the MuSK-BMP pathway plays a role in SC quiescence in older mice, we next assessed SC number and myofiber diameter at 5 months of age. Both measures showed striking differences between genotypes at this age. SC density in the 5-month-old TA was reduced by almost half in ΔIg3-MuSK compared to WT mice (19±2.1 and 10.6±1.4mm^2^ in WT vs ΔIg3-MuSK, respectively; n=9-10, p=0.004; **Fig. 2a, c**). On the other hand, myofiber diameter in ΔIg3-MuSK muscle was increased by ∼10% compared to WT (37 ±0.8µm and 41 ± 1.0µm in WT vs ΔIg3-MuSK, respectively; n=8, p=0.006; **Fig. 2a, b**).

### Grip strength is increased in 5-month-old ΔIg3-MuSK mice

We next asked whether the increase in myofiber size observed in 5-month-old mice was associated with improved muscle function. We performed grip strength analysis of ΔIg3-MuSK male mice at 3- and 5- months of age. As shown in **Fig. 2**, grip strength is comparable between ΔIg3-MuSK and WT mice at 3 months. In contrast, at 5 months, male ΔIg3-MuSK mice display increased grip strength performance (**Fig. 2d**). This increase in grip strength was observed in 3 cohorts of male mice (Supp. Fig. 2). We also tested grip strength in one cohort of female mice. Although the average grip strength was higher in 5 month old ΔIg3-MuSK females compared to controls, this effect did not reach statistical significance (**Supp. Fig. 2**). Thus, muscle function is improved in 5- month-old male ΔIg3-MuSK animals.

### Myonuclei numbers and myofiber lengths are increased in ΔIg3-MuSK EDL muscles at 5 months of age

The observation that SC numbers are decreased while myofiber diameter is increased in 5- month-old ΔIg3-MuSK TA muscle suggested that SCs in these adult muscles became activated and fused into the myofiber, thus contributing to its growth. To test this idea directly we counted the number of myonuclei in isolated myofibers from 5-month-old ΔIg3-MuSK and WT EDL muscles, an accessible and readily quantifiable model for such measurements. As shown in **Fig. 3a, b**, the number of myonuclei/myofiber was increased in ΔIg3-MuSK EDL muscle. In addition, the ΔIg3-MuSK myofibers were also longer **(Fig. 3a, c)**. Importantly, the proportion of myonuclei per myofiber length is equivalent at both ages, suggesting that the size of the myonuclear domain is conserved (**Fig. 3d**). Taken together, these findings indicate that between 3 and 5 months of age in ΔIg3-MuSK muscle SCs become activated, divide, and fuse into the myofiber - resulting in increased length. Further, the conserved proportion of myonuclei/myofiber length indicates that this growth increase scales with the addition of the myonuclei to the myofiber.

**Figure 3.**
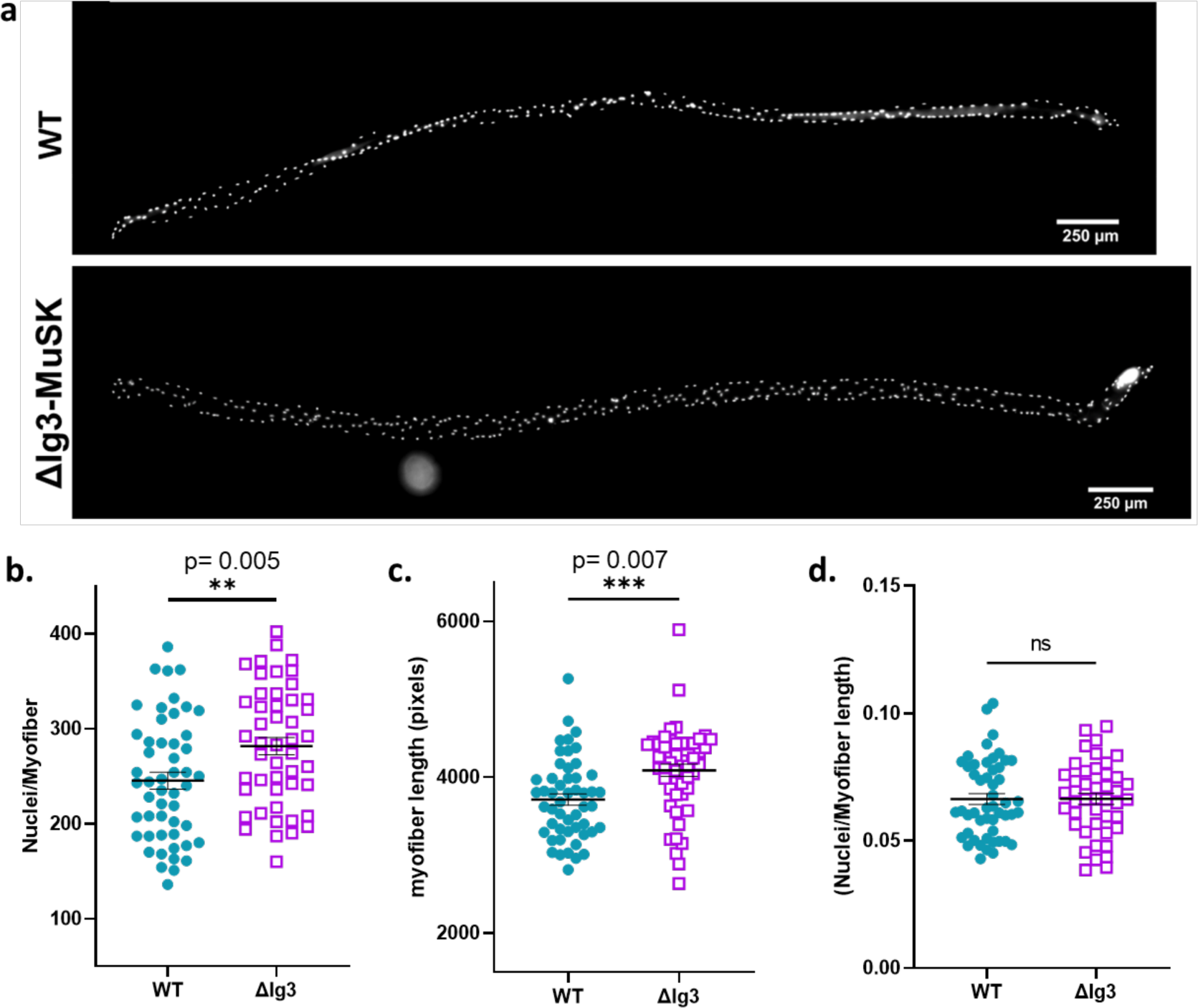
Increased myofiber myonuclei number and length in 5-month-old ΔIg3-MuSK EDL. (**a**) Representative DAPI-stained images of isolated WT and ΔIg3-MuSK EDL myofibers from 5 mo. old mice. (**b**) Note that myonuclear number/myofiber is increased ∼13% in ΔIg3-MuSK compared to WT (245 ±9.0 and 282±9.0 in WT and ΔIg3-MuSK myofibers, respectively; n=5 male mice, 7-10 myofibers per mouse; p=0.005; unpaired t-test). (**c**) 5-month-old ΔIg3-MuSK myofibers length is increased ∼10% compared to WT (3714±72 and 4089±79 in WT and ΔIg3-MuSK myofibers, respectively; n=5 male mice, 7-10 myofibers per mouse; t-test, p=0.0007). (**d**)The numbers of nuclei per myofiber length was comparable in ΔIg3-MuSK and WT myofibers (n=5 male mice, 7-10 myofibers per mouse; unpaired t-test, p=0.99)

### Muscle regeneration and restoration of SC numbers following injury to ΔIg3-MuSK muscle

We next queried the role of ΔIg3-MuSK in muscle regeneration, a chief function of adult SCs. TA muscles from 5-month-old WT and ΔIg3-MuSK TA were injured on day 0 by BaCl_2_ injection and muscles were harvested at 3-, 5-, 7-, 14-, 22-, and 29-days post-injury (dpi; **Fig. 4**). We assessed muscle morphology, myofiber diameter, and SC density at each timepoint. The ΔIg3-MuSK muscle regenerated efficiently, with full restoration of muscle structure achieved by 22 dpi, which was comparable to WT (**Fig. 4b, c, e**). The regeneration rate was increased as judged by histoarchitecture (**Fig. 4b, c**), increased Pax7+ density at 5 dpi (**Fig. 4d**) and, in the male animals, increased myofiber diameter at 7 dpi (**Fig. 4e**). The increased Pax7+ cells at 5 dpi and the increased myofiber size at 7dpi indicate that the early stages of muscle regeneration are accelerated in ΔIg3-MuSK mice.

**Figure 4.**
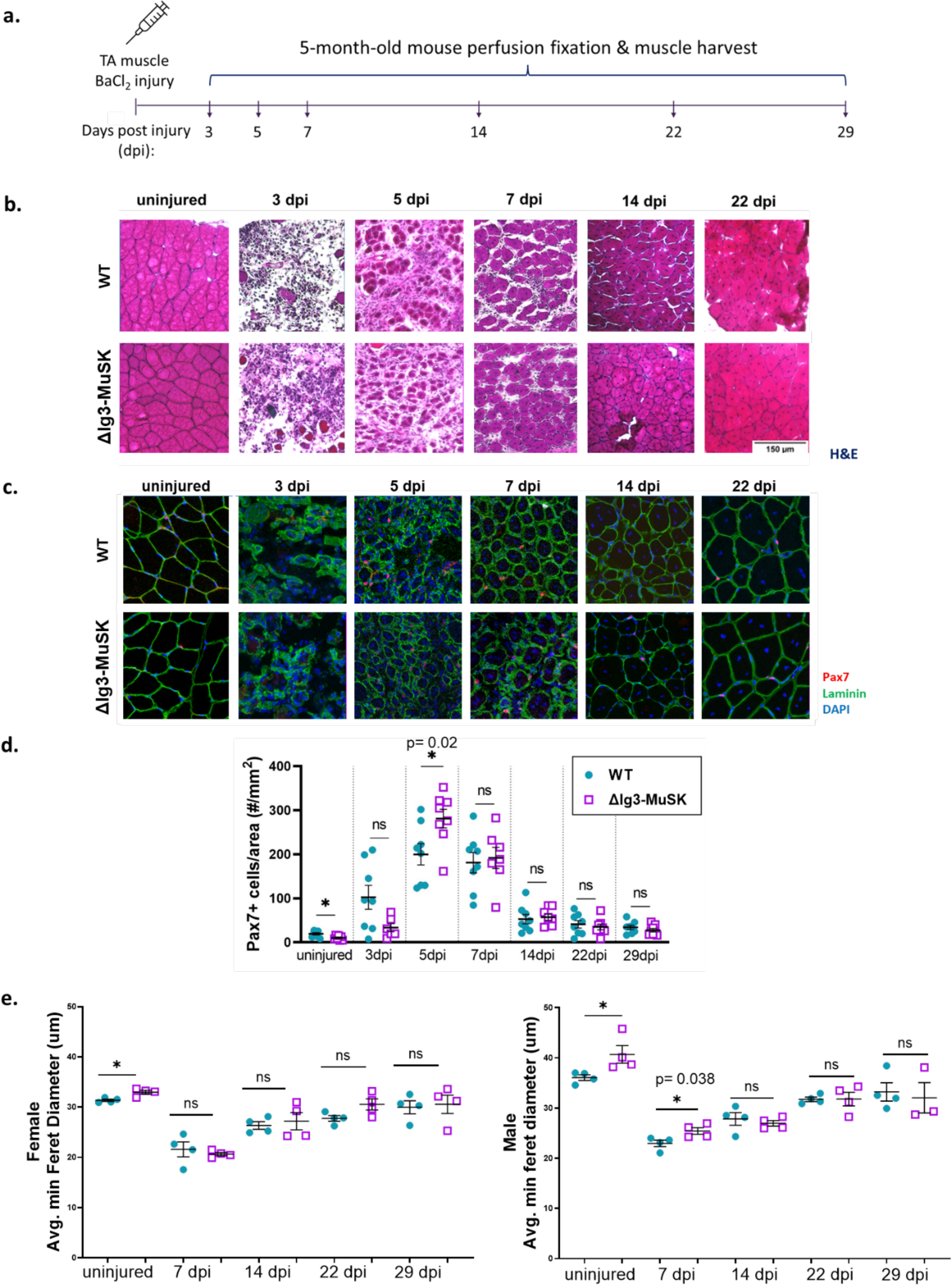
Efficient, accelerated regeneration in 5-month-old ΔIg3-MuSK muscle. (**a**) Experimental design. WT and ΔIg3-MuSK TA muscles were injected with BaCl_2_ at day 0 and muscle harvested at the indicated time points. (**b**) Hematoxylin and eosin stain of uninjured and regenerating ΔIg3-MuSK and WT male TA muscle. (**c.**) IHC images of uninjured and regenerating ΔIg3-MuSK and WT male TA muscle used to quantify Pax7+ cells and myofiber size; laminin (green), Pax7 (red), DAPI (blue). (**d**) The number of Pax7+ cells is reduced in resting control ΔIg3-MuSK muscle at 5 months (see also Fig. 2a, c). However, during regeneration the number of Pax7+ cells in ΔIg3-MuSK muscle is increased relative to WT at 5 dpi (n=8 mice (4 male, 4 female; 5dpi t-test p= 0.023). (**e**) Analysis of myofiber size in females (left) and males (right) reveals increase in minimum Feret diameter of ΔIg3-MuSK myofibers in uninjured mice (see also fig. 2b). Following injury, an increase in myofiber size was observed at 7 dpi in ΔIg3-MuSK males (n=4 mice/sex, t-test, Male 7dpi p= 0.038). No significant differences in Feret diameter were observed in regenerating female WT compared to ΔIg3-MuSK muscle.

Notably, the myofiber diameters at the completion of regeneration (22 dpi) were comparable in both genotypes (**Fig. 4e)**. Moreover, the density of SCs in the regenerated muscle (22 and 29 dpi) was indistinguishable in WT and ΔIg3-MuSK (**Fig. 4d**). This comparable restoration of myofiber size and SC density indicates that the intrinsic regulatory mechanisms governing SC dynamics during regeneration, including stemness and regenerative capacity, are maintained in ΔIg3-MuSK muscle.

### MuSK expressed in adult SCs governs quiescence and myofiber hypertrophy

The studies presented above utilized mice with constitutive (germline) knock-in of the ΔIg3-MuSK allele. However, MuSK is also highly expressed by neuromuscular junction (NMJ) myonuclei, and at lower levels in subsets of myonuclei and interstitial cells(Jaime et al., 2024; Petrany et al., 2020; Zeng et al., 2022; Zhang et al., 2021). Thus, there were two key questions outstanding: 1) Is the function of the MuSK-BMP pathway in SC dynamics cell autonomous; and 2) are there developmental events regulated by this pathway that might contribute to its role in adult SCs? To address both these points we utilized tamoxifen-inducible, conditional *Pax7^CreERT2^;R26R^LSL-tdTomato^;MuSK-Ig3^loxP/loxP^* mice. These mice license the expression of ΔIg3-MuSK only in cells expressing Pax7 and at the age determined by the tamoxifen administration. In addition, it marks them with the TdTomato reporter. As a control, we used *Pax7^CreERT2^;R26R^LSL-tdTomato^*, in which tamoxifen injection induces TdTomato expression in Pax7+ cells, but the MuSK allele is unaffected. We induced Cre-recombination using tamoxifen administered at 3 months of age and then harvested the TA muscles at 5 months of age (**Fig. 5a**). PCR analysis of RNA isolated from pre-fixed, FACS-sorted SCs confirmed that the Cre-recombination of the floxed Ig3 domain of MuSK was effective (**Supp. Fig. 3a, b**). As shown in **Fig. 5**, Cre-mediated expression of ΔIg3-MuSK in SCs at 3 months resulted in an ∼9% increase in both muscle weight and myofiber diameter when assessed at 5 months of age (0.035±0.0007 and 0.038±0.0007 g, n=5-8 mice/group, p=0.007; and 36.0±0.78 and 39.3±1.2 µ, n=5-7 mice/group, p=0.037 for weights and diameters, respectively; **Fig. 5**). Finally, we also counted the number of myonuclei per myofiber in EDL muscles from the same animals (see Methods). **Fig. 5e** shows that there was an ∼22% increase in the myonuclear number in the *Pax7^CreERT2^;R26R^LSL-tdTomato^;MuSK-Ig3^loxP/loxP^* compared to controls (*Pax7^CreERT2^;R26R^LSL-tdTomato^*; 234.2±9.8 and 286. ±16.2, respectively, t-test, p=0.008; n=3 per genotype). The increased TA weight, myofiber size and myonuclear number observed in these experiments indicates that MuSK acts in a SC-autonomous manner and that the expression of MuSK in the adult SC is sufficient for this role.

**Figure 5.**
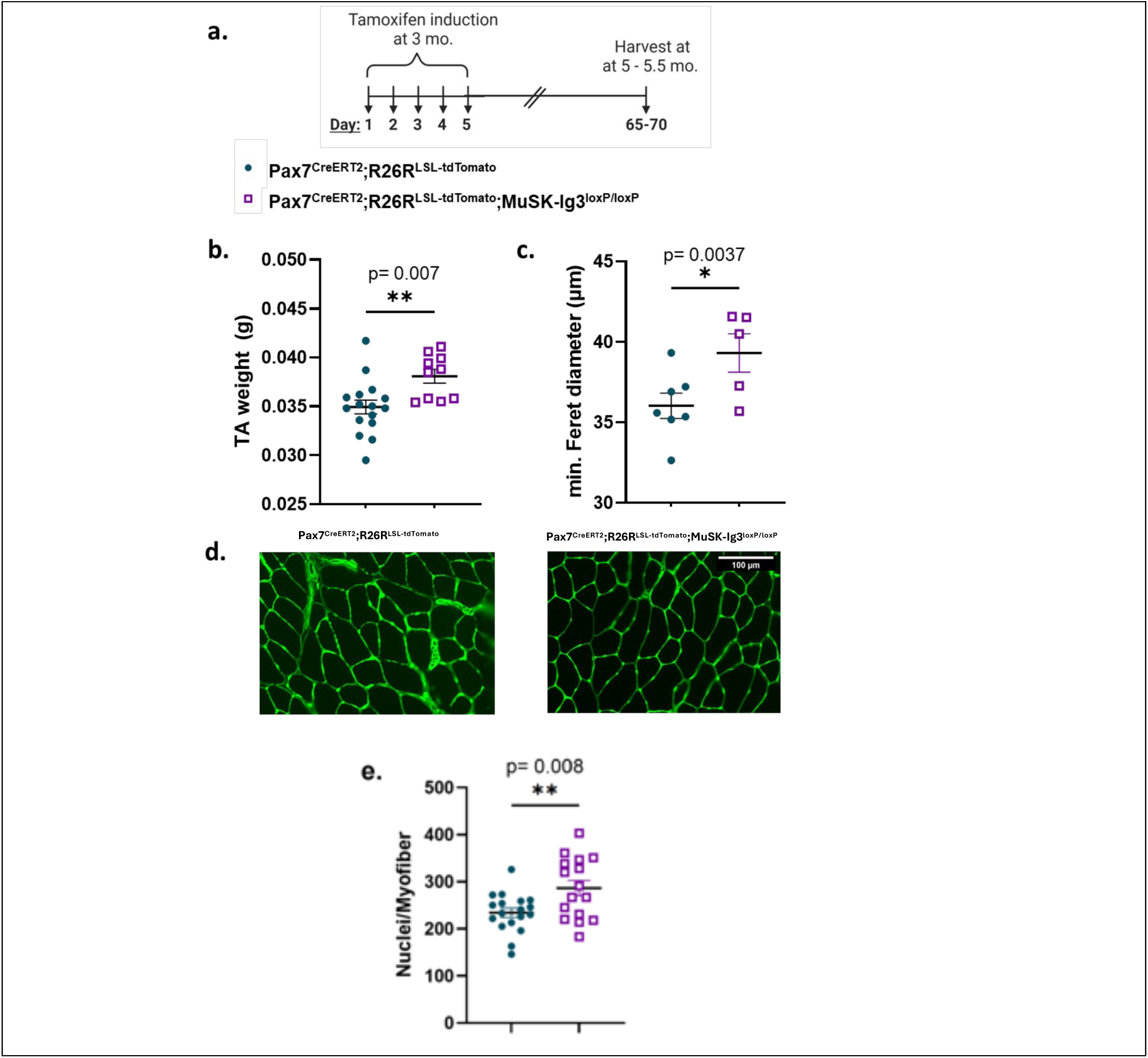
Expression of ΔIg3-MuSK in SCs from 3 months of age is sufficient to mediate increased TA muscle weight and myofiber size at 5 months. (**a.**) Experimental design. At 3 months of age tamoxifen was administered for 5 consecutive days to *Pax7^CreERT2^;R26R^LSL-tdTomato^;MuSK-Ig3^loxP/loxP^*(test group, purple) and *Pax7^CreERT2^;R26R^LSL-tdTomato^* (control group, dark teal) mice, which were harvested 65-70 days later at ∼5 months of age. (**b.**) TA muscle weights were increased ∼9% in *Pax7^CreERT2^;R26R^LSL-tdTomato^;MuSK-Ig3^loxP/loxP^*mice (n=5-8 mice/group; 2 TAs per mouse), (0.035±0.0007 and 0.038±0.0007 g; t-test, p=0.007). (**c.**) Myofiber cross-sectional minimum Feret diameter was also increased (∼9%) in *Pax7^CreERT2^;R26R^LSL-tdTomato^;MuSK-Ig3^loxP/loxP^* mice (n=5-7 mice/group; 36.0±0.78 and 39.3±1.2; t-test, p=0.037). (**d.)** Representative images of *Pax7^CreERT2^;R26R^LSL-tdTomato^* (control, left) and *Pax7^CreERT2^;R26R^LSL-tdTomato^;MuSK-Ig3^loxP/loxP^*(test, right) representative IHC images of anti-laminin (green) used to measure myofiber size. (**e.**) Nuclei number per EDL myofiber was increased (∼22%) in test compared to controls (234.2±9.8 and 286. ±16.2, respectively, t-test, p=0.008).

### SCs are activated in uninjured ΔIg3-MuSK muscle

The results presented above indicate that the MuSK-BMP pathway is important for maintaining SC quiescence in resting adult muscle, and that perturbation of this pathway leads to the activation of these stem cells. To provide direct evidence for such activation and to gain insight to the mechanistic basis of this state change we interrogated the gene expression of FACs-sorted SCs from uninjured 5-month-old WT and ΔIg3-MuSK muscle (n= 6 animals/genotype). To preclude artefactual activation during cell isolation, animals were perfusion-fixed prior to harvest. We then used the NanoString nCounter Stem Cell Characterization Panel to assess gene expression in WT and ΔIg3-MuSK SCs. This panel includes 770 genes involved in multiple stem cell types including stem-cell regulatory signaling pathways, epigenetics, metabolism, differentiation signaling, and lineage specification. Hierarchical clustering revealed significant changes (p < 0.05) in the expression of 21 of the genes in this panel. Nineteen of the differentially expressed genes (DEGs) were upregulated including TBLX1, HDAC2, PPP2R2A, SMAD4, CAP2, CREBBP, MDH1, NCOR1, RRAGC and CUL1; while two genes, XIST and CACYBP, were downregulated (**Fig. 6a, b**).

**Figure 6.**
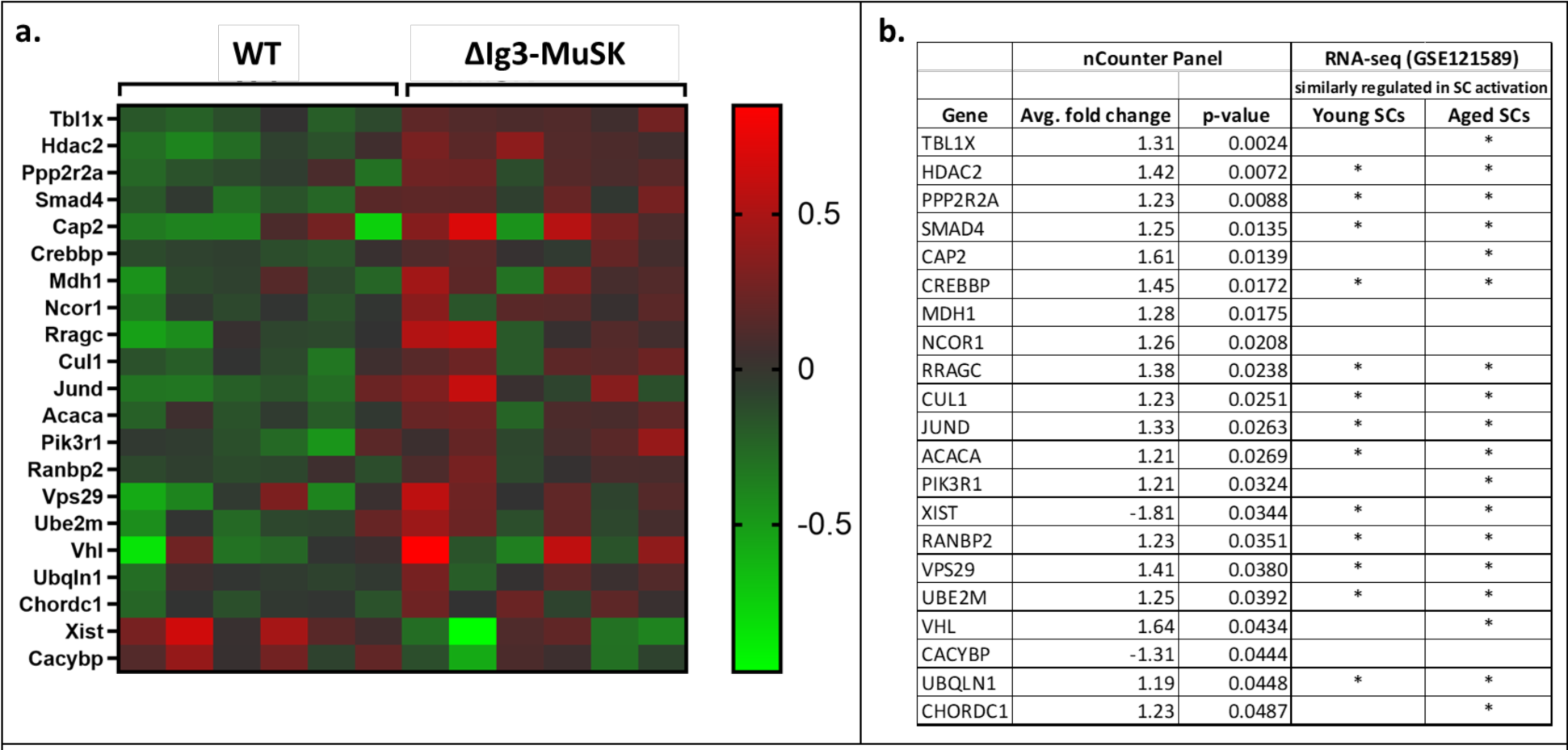
SCs are activated in uninjured ΔIg3-MuSK mice. (**a**) Heatmap of 21 differentially expressed genes between the two genotypes detected by nCounter analysis using the Stem Cell Characterization Panel. **(b)** Comparison of the DEGs in WT and ΔIg3-MuSK SCs detected by nCounter analysis with the DEGs detected in SCs from either ‘young’ or ‘aged’ animals by RNAseq analysis during early damage-induced activation (uninjured vs. 1 dpi; GSE121589). “*” indicates whether each gene was significantly upregulated or downregulated in the respective data sets.

We next asked whether the set of DEG SC genes detected in uninjured ΔIg3-MuSK were characteristic of an activated state. We compared the hits in the ΔIg3-MuSK nCounter set to a published RNAseq dataset of SC DEGs from resting and injured (1 dpi) WT muscle from young (2-3 months) and aged (20-22 months) muscle (GSE121589; Shcherbina et al., 2020). Our analysis revealed striking similarities in the sets of SC DEGs from uninjured ΔIg3-MuSK and damaged WT muscles. As shown in **Fig. 6a, b,** 17 of the 19 upregulated genes in uninjured ΔIg3-MuSK SCs are also upregulated in SCs from old and/or young activated WT SCs. XIST, one of the two genes downregulated in ΔIg3-MuSK SCs, was also reduced in both young and aged SCs from damaged WT muscle. This concordance of gene expression patterns indicates that ΔIg3-MuSK SCs in 5-month-old uninjured muscle are in a state of early activation.

## DISCUSSION

Our results establish that the BMP co-receptor MuSK regulates SC quiescence in adult muscle in a cell-autonomous fashion. Importantly, perturbation of this pathway leads to SC activation, proliferation, and fusion - culminating in increased muscle weight and grip strength. Stemness is maintained (**Fig. 7**). Below we discuss these novel features of adult SC quiescence and MuSK function as well as their implications for therapeutic development for muscle wasting conditions.

**Figure 7.**
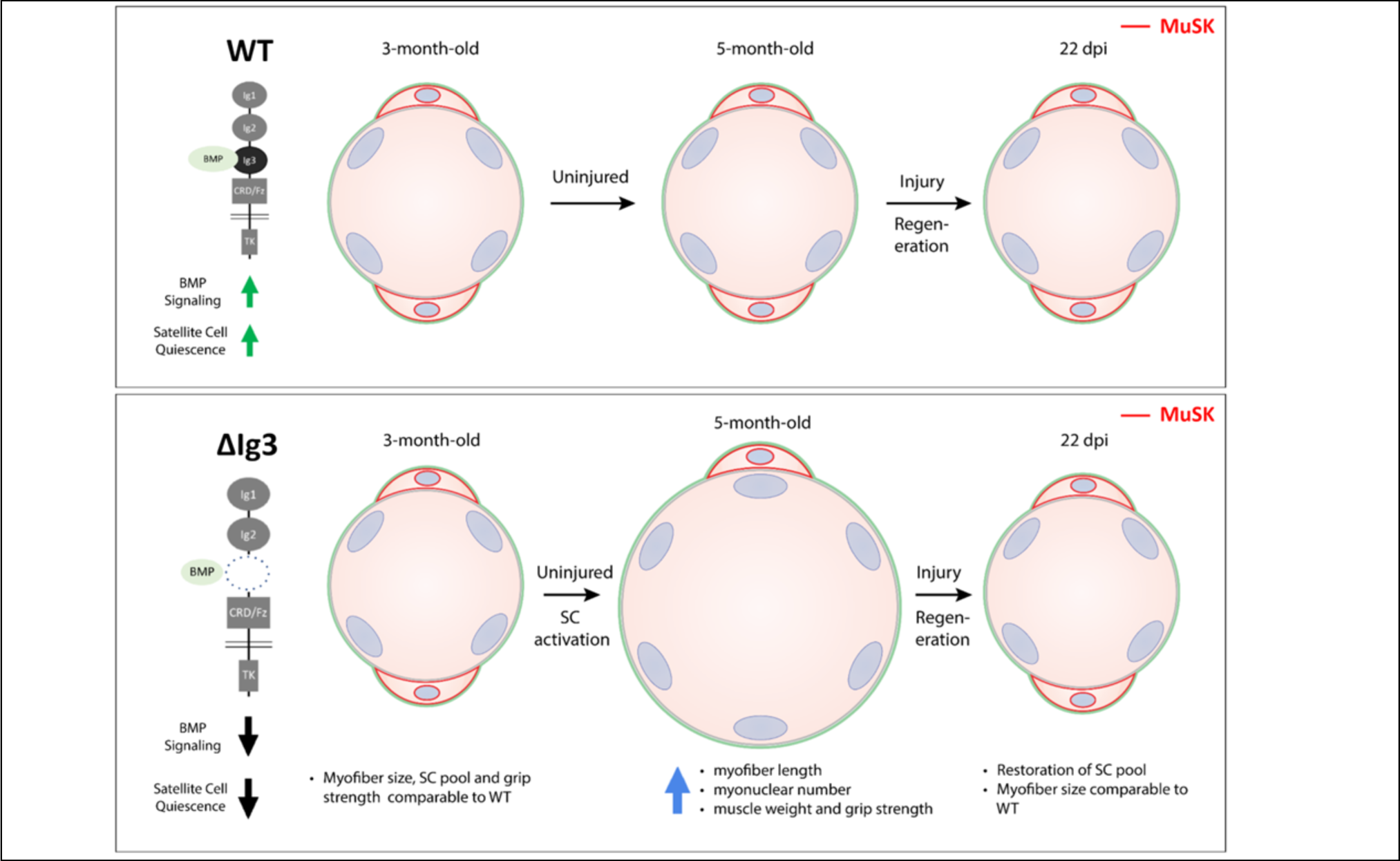
A model for the function of the MuSK-BMP pathway in adult SCs. MuSK is expressed in both WT and ΔIg3-MuSK SCs (*red*) where it functions in a cell autonomous manner. **Top.** In wildtype the MuSK Ig3 domain mediates high affinity BMP binding and MuSK-BMP signaling, maintaining SC quiescence and myofiber size. **Bottom.** In ΔIg3-MuSK, BMP binding and signaling are reduced, disrupting SC quiescence. Between 3- and 5- months of age SCs expressing ΔIg3-MuSK are activated in uninjured adult muscle and divide and fuse into the myofiber, resulting in increased myofiber size, myonuclear number and grip strength. SCs in 5-month-old ΔIg3-MuSK muscle maintain stemness and can support regeneration and reestablishment of the SC pool following injury.

MuSK regulates SC quiescence in a cell autonomous manner. MuSK transcript and protein are expressed in SCs (**Fig. 1**). This finding is in accord with snRNAseq studies showing MuSK transcript in quiescent SCs(Petrany et al., 2020; Shcherbina et al., 2020; Tabula Muris Consortium et al., 2018; Zeng et al., 2022). Our results also show that quiescent SCs express ‘full length’ MuSK transcript encoding the Ig3 domain (**Fig. 1; Supp. Fig. 1**). This point is noteworthy since the Ig3 domain can be endogenously spliced(Garcia-Osta et al., 2006; Hesser et al., 1999). Finally, conditional expression of ΔIg3-MuSK in SCs at 3 months of age is sufficient to phenocopy the increase in myofiber size and myonuclear numbers observed in germ line mutants. Thus, MuSK containing its Ig3 domain is necessary for maintaining adult SC quiescence.

Several lines of evidence indicate that MuSK in SCs functions through its role as a BMP co-receptor, and not as an agrin-LRP4 receptor. Biochemical studies show that MuSK binds BMPs with high affinity in a manner requiring its Ig3 domain(Yilmaz et al., 2016). Quiescent SCs express BMP signaling machinery, including BMP4 as well as BMPR1a, which also binds to MuSK(Petrany et al., 2020; Shcherbina et al., 2020; Yilmaz et al., 2016; Zeng et al., 2022). Cultured ΔIg3-MuSK myoblasts have reduced BMP4 responses as assessed by pSMAD signaling and transcriptional output(Jaime et al., 2024). In contrast, the data does not support a role for MuSK acting as an agrin-LRP4 receptor in SCs: 1) Agrin-induced AChR clustering, which is mediated by its interaction with LRP4, is comparable in ΔIg3-MuSK and WT myotubes(Jaime et al., 2024); 2) NMJ formation, maintenance, AChR density and cholinergic signaling is comparable in WT and ΔIg3-MuSK muscle. Interestingly, NaV1.4 density and muscle excitability is reduced, suggesting that the MuSK-BMP pathway plays distinct roles from agrin-LRP4 at the NMJ (Fish et al., 2023); ( 3) MuSK’s function as a BMP co-receptor requires neither its TK activity nor the juxtamembrane tyrosine critical for DOK7 binding – features necessary for agrin-LRP4 activity(Jaime et al., 2024). 4) The MuSK Ig3 domain is dispensable for agrin-LRP binding and signaling, which is mediated by its Ig1 domain(Hesser et al., 1999; Stiegler et al., 2006; Yilmaz et al., 2016; Zong et al., 2012). To our knowledge this is the first report implicating BMP signaling in adult SC quiescence. It will be of interest to test whether MuSK also plays a role in the BMP-mediated quiescence regulation in other settings, such as neural stems cells(Díaz-Moreno et al., 2018; Mira et al., 2010; Tunc-Ozcan et al., 2021). Finally, the deployment of MuSK, which has a highly restricted cell type and tissue distribution, as a co-receptor could be an evolutionary solution to the general problem of conferring context selectivity to the near-universally expressed BMPs and their core signaling machinery(Hogan, 1996; Massagué, 2012).

Our results show that adult ΔIg3-MuSK satellite cells are activated in uninjured muscle. First, SCs density decreases by ∼50% from 3- to 5- months of age in the ΔIg3-MuSK TA. Second, transcriptomic analysis shows that the expression profile of SCs from uninjured adult muscle is remarkably similar to that observed in WT muscle 1 day post injury (**Fig. 6**;(Shcherbina et al., 2020). Third, the accelerated muscle regeneration following injury observed in ΔIg3-MuSK mice mirrors that observed in other settings where SCs are activated, such as deletion of the myofiber-derived quiescence factor Wnt4 or injury to the contralateral muscle(Eliazer et al., 2019; Rodgers et al., 2014). This accelerated regeneration likely reflects a more rapid re-entry into the cell cycle(Siegel et al., 2011).

MuSK plays a selective role in maintaining adult SC quiescence. SCs are highly dynamic and assume multiple critical states including, quiescence, activation, proliferation, fusion into myofibers, and re-entry into quiescence. Adult SCs expressing ΔIg3-MuSK SC break quiescence and become activated, but their ability to proliferate and fuse into myofibers is preserved. By 5- months of age the ΔIg3-MuSK EDL has increased length, adding an average of 40 myonuclei/myofiber compared to WT (**Fig. 3a, c**). As EDL myofibers contain only ∼7 SCs each, the activated SCs must undergo at least 3 rounds of cell division over this time span, followed by fusion into the myofiber. These traits distinguish the MuSK-BMP pathway from many other mechanisms mediating quiescence. For example, manipulation of Notch, Foxo3 or Pten signaling breaks quiescence, but the SC pool is depleted and self-renewal is impaired(García-Prat et al., 2020; Gioftsidi et al., 2022; Yue et al., 2017). Disruption of the M- and N- cadherins or Wnt4a from the myofiber results in expansion of the SC pool, but no fusion or changes in myofiber diameter were reported(Eliazer et al., 2019; Goel et al., 2017). Finally, interfering with the ColV-Calcitonin receptor pathway in SCs leads to spontaneous differentiation and fusion, but SCs were fewer and myofibers smaller following damage(Baghdadi et al., 2018; Zhang et al., 2019). Taken together, these results indicate that MuSK-BMP signaling restrains SC activation, but is dispensable for proliferation and productive fusion into uninjured myofibers. Notably, all three of these activities are critical for SCs to contribute to the muscle growth and improved strength such as observed here. Importantly, following injury myofiber size and SC complement are restored to WT levels, indicating that MuSK-BMP signaling is not required for regeneration, to maintain stemness, or for re-entry into the SC niche. Moreover, the normal fiber size and SC complement observed in germ line mutants at 3 months indicates that the MuSK-BMP pathway does not play a discernable role in SCs in developing and young mice, a conclusion supported by the results with conditional expression of ΔIg3-MuSK in SCs starting at 3 months (Fig. 5). Finally, the adult-selective role of the MuSK-BMP pathway is distinct from myostatin (GDF8), which has profound effects on SCs proliferation and hyperplasia during early postnatal development, but does not regulate SC quiescence in adult muscle(Lee, 2022; Murphy et al., 2010).

The MuSK-BMP pathway represents an attractive therapeutic target to combat muscle wasting and promote regeneration. Muscle size and function are compromised in settings including muscular dystrophies, cancer cachexia, disuse atrophy, age-related sarcopenia, and GLP-1 agonist use (Bikou et al., 2024; Chamberlain and Olwin, 2010; Ham et al., 2022; Shavlakadze et al., 2019). Muscle growth is limited by myonuclear complement(Cramer et al., 2020). SCs are present in these situations and represent a potential source of new myonuclei. However, the SCs in disease and aging are often in deep quiescence due to degradation of the niche and the accumulation of negative signals countering activation(Brunet et al., 2023; Cosgrove et al., 2014; Kimmel et al., 2020; Liu et al., 2017). Notably, MuSK levels are increased in mouse models of denervation and Myasthenia Gravis as well as in aging humans, suggesting that full length MuSK may generate one such negative signal(Bowen et al., 1998; Lehallier et al., 2019; Mori et al., 2023). Targeting this pathway could also promote longevity and healthy aging. Reducing MuSK-BMP signaling to promote SC activation is an attractive, highly specific target given the restricted tissue distribution of MuSK(Tabula Muris Consortium et al., 2018), combined with the precise modulation of the Ig3 domain as demonstrated genetically in this report. Pharmacological targeting could be achieved, for example, by splice-modulating antisense oligonucleotides or antibodies directed against the MuSK Ig3 domain. The normal life span of mice with germ line expression of ΔIg3-MuSK suggests a favorable safety profile for such therapeutic interventions.

## MATERIALS AND METHODS

### Mice

Mice were bred and housed at the Brown University Animal Care Facility (ACF). Animals were group-housed in cages of up to five animals. All procedures were approved by the Institutional Animal Care and Use Committee (IACUC) at Brown University and done in accordance with institutional guidelines for animal care.

Germ line MuSK^ΔIg3/ΔIg3^ mice, referred to throughout this paper as ‘ΔIg3-MuSK’ mice, were generated by Crispr-mediated removal of exons 6 and 7 of MuSK, which encode the Ig3 domain and were on the C57Bl/6 background. Details of the production and characterization of this model have been reported(Jaime et al., 2024).

For conditional mutations we generated a MuSK-Ig3^loxP/loxP^ mouse using CRISPR/Cas9, in which the exons encoding the Ig3 domain of MuSK (exons 6 and 7) are flanked with loxP sites. Tamoxifen-inducible *Pax7^CreERT2^;R26R^LSL-tdTomato^*mice were provided by Bradley Olwin(Pawlikowski et al., 2015). These lines were crossed to generate *Pax7^CreERT2^;R26R^LSL-tdTomato^;MuSK-Ig3^loxP/loxP^*mice. Recombination was induced with five daily intraperitoneal (IP) injections of tamoxifen (Sigma-Aldrich) in corn oil dosed at 0.1 mg tamoxifen/ g mouse weight. Mice were treated with tamoxifen at 3 months of age to induce Cre-recombination (**Supp. Fig. 2**).

### Muscle Harvest

For Pax7 antibody staining muscles were harvested from perfusion-fixed animals. Briefly, animals were anesthetized using 2,2,2-Tribromoethanol (250 mg/kg, Sigma Aldrich) followed by perfusion with 20mL of 4% PFA in PBS prior to TA harvest. Muscles were dehydrated in 30% sucrose in PBS overnight, frozen in OCT using liquid nitrogen, and stored at −80°C. In other experiments mice were euthanized with CO_2_ followed by cervical dislocation. TA muscles were then flash frozen using freezing isopentane and stored at −80°C.

### Immunohistochemistry

Frozen sections (10µm) were obtained on a Leica Cryostat. Sections from flash frozen muscle were post-fixed in 4% paraformaldehyde (PFA) for 10 minutes at room temperature. Tissue sections and isolated myofibers were permeabilized with 0.25% Triton-X100 (Sigma-Aldrich) in PBS followed by blocking with 2% bovine serum albumin (Sigma-Aldrich) and 5% goat serum (Thermo Fisher) in PBS. Sections and myofibers that were stained with anti-Pax7 antibody (DSHB) also included a block with 1:50 Mouse on Mouse Ig Blocking Reagent (MKB-2213, Vector Labs). Incubation with primary antibody was at 4°C overnight followed by incubation with secondary antibody at room temperature for 1 hr. To visualize EdU incorporation, the Click-it Plus EdU Cell Proliferation kin, Alexa flour-488 kit (C10637, Thermo-Fisher) was used following the manufacturer’s instructions. Primary antibodies were 1:20 anti-Pax7 (Developmental Studies Hybridoma Bank), 1:1000 rabbit anti-laminin (L9393, Sigma-Aldrich), MuSK myasthenia gravis (MG) patient IgG4 fraction was generously provided by M. Huijbers(Huijbers et al., 2013). Secondary antibodies against IgG1 or IgG of the appropriate species were conjugated to Alexa-488, Alexa-555, or Alexa-647 (Invitrogen) and used at 1:1000. Myofibers were incubated with 1µg/mL DAPI (D1306, Invitrogen) for 5 minutes at room temperature and then mounted in Mowiol supplemented with DABCO (Sigma-Aldrich) as an anti-fade agent. Sections were mounted with Vectashield+DAPI as an anti-fade agent (Vector Laboratories).

### Immunofluorescence

Immunofluorescence imaging was performed using a Nikon E800, a Nikon Ti2-E Widefield, or a Zeiss LSM 800 Confocal Microscope. Data acquisition was performed using the Zen software (blue edition, Zeiss). Images used for Pax7 counting and myofiber size analysis were collected and scored blinded.

### Myofiber cross sectional size

ΔIg3-MuSK mouse muscles used for myofiber cross sectional size analysis were from perfusion fixed animals and *Pax7^CreERT2^;R26R^LSL-tdTomato^;MuSK-Ig3^loxP/loxP^* muscles used for myofiber cross sectional size analysis were from flash frozen muscle. Images of TA muscle sections immunostained with anti-laminin antibody (L9393, Sigma-Aldrich) and captured using a Nikon E800 fluorescent microscope or with a Zeiss LSM 800 Confocal Microscope with a 20X objective (images within the same experiment were captured using the same microscope). Images were processed using MyoVision software to determine minimum Feret diameter measurements for each myofiber(Briguet et al., 2004; Wen et al., 2018). Output images were manually examined, and any misidentified fibers were excluded, 650-1800 fibers examined per mouse. Representative images were acquired with a Zeiss LSM 800 Confocal Laser Scanning Microscope with a 20x objective. The average myofiber min Feret diameter for each mouse was used for statistical analysis using an unpaired t-test in GraphPad Prism 9. Analysis was done blinded.

### Intact myofiber isolation

Extensor digitorum longus (EDL) muscle was dissected and enzymatically digested in 400-U/mL collagenase II (Worthington) at 37°C for 1.5 hours. Collagenase was inactivated by the addition of DMEM supplemented with 15% horse serum. Flame-polished Pasteur pipettes were used to agitate and separate individual myofibers, which were then immediately fixed in 4% paraformaldehyde for 10 minutes, followed by 3 washes of PBS prior to any immunostaining.

### Intact myofiber myonuclei and length quantification

After immunostaining myofibers (see “Immunohistochemistry” methods), z-stacked tiled images were taken of whole myofibers with a Nikon Ti2-E Widefield Fluorescence Microscope using Zen software (blue edition, Zeiss). The automated measurement function was used to quantify all DAPI stained nuclei as well as total myofiber length.

### Grip Strength

Grip strength analysis was done to assess front-limb muscle strength of WT and ΔIg3-MuSK male and female mice at 3 and 5.5 months of age. Mice were lifted by their tail and guided to use both front paws to grasp the pull-bar assembly connected to the grip strength meter (Columbus Instruments). The mouse was pulled away from the sensor until their grip on the bar was broken and the peak amount of force in kilograms per unit force (KGF) was recorded. Five trials were performed sequentially per animal and then averaged together and normalized to mouse weight. Analysis was done blinded to genotype.

### Muscle injury & EdU delivery

Briefly, 5-month-old mice were anesthetized with isoflurane (induced with 4% isoflurane, maintained at ∼1.5% isoflurane and 400 mL O2 per min) and placed on a temperature-controlled platform to maintain core body temperature between 35°C and 37°C. Injury to the left TA muscle was induced by first piercing the muscle with a 28-gauge needle ∼15 times followed by injection of 50*μ*L of 1.2% BaCl2 in PBS. To relieve pain, mice were treated with 0.5mg/mL of Buprenorphine Sustained-Release (Bup-SR) at the time of injury. For EdU labeling, mice received 50mg/kg 5- ethynyl-2’-deoxyuridine (EdU, Thermo Fisher) in PBS by intraperitoneal injection 24 hr before harvest. Injured muscle was harvested at 3-, 5-, 7-, 14-, 22-, and 29-days post injury (dpi). Control muscle from uninjured mice was also collected.

### Satellite cell isolation and FACS sorting

To prevent artificial activation of quiescent SCs during isolation, mice were perfusion fixed with 0.5% PFA and then muscle was digested with collagenase II (Worthington) as described(Yue et al., 2020; Yue and Cheung, 2020). Pooled hind limb muscles were used for analysis of SCs from resting muscle and pooled TA muscles were used for analysis of SCs from regenerating muscle 5 dpi. The resulting cell suspension was blocked with DMEM + 15% horse serum (Cytiva sh30074) for 10 min at 4°C with rocking. Cells were then incubated for 45 min at 4°C with rocking with 1:250 of each of the following antibodies: Negative selection markers: BV421 Rat anti-mouse anti-CD31 (562939, BD Biosciences), and BV421 Rat anti-mouse CD45 (563890, BD Biosciences), BV421 Rat anti-Mouse Ly-6aA/E (562729, BD Biosciences); positive selection marker: Mouse anti-integrin alpha 7 Alexa fluor 647 (FAB3518R, R&D Systems). Cells were rinsed, resuspended in PBS and sorted using a BD FACSAria IIu cell sorter equipped with 355nm, 407nm, 488nm, 561nm, and 633nm lasers. The machine was optimized for purity and viability. UltaComp eBeads were used for calibration (01-222-41, Thermo Scientific). Total RNA was isolated from FACS-sorted SCs using a miRNeasy FFPE Kit (Qiagen) as described(Yue and Cheung, 2020).

### Nanostring nCounter assay

RNA from SCs of 5 - 5.5-month-old resting muscle (n=6/genotype) was isolated as outlined above and samples from all mice were kept separate. Gene expression was analyzed with a Nanostring nCounter Stem Cell Characterization panel (NanoString Technologies). Multiplex gene expression with 770 genes was performed by measurement of the abundance of each mRNA transcript of interest using multiplexed hybridization and digital readouts of fluorescent bar-coded probes that are hybridized to each transcript. Hybridization of samples was performed, and the obtained products were run according to the manufacturer’s instructions. Data were collected on the nCounter Sprint profiler and further analyzed by _ROSA_LIND® (https://rosalind.bio/), with a HyperScale architecture developed by ROSALIND, Inc. (San Diego, CA). Read Distribution percentages, violin plots, identity heatmaps, and sample MDS plots were generated as part of the QC step.

ROSALIND® follows the nCounter® Advanced Analysis protocol of dividing counts within a lane by the geometric mean of the normalizer probes from the same lane. Housekeeping probes to be used for normalization are selected based on the geNorm algorithm as implemented in the NormqPCR R library(Perkins et al., 2012). Fold changes and p values are calculated using the fast method as described in the nCounter® Advanced Analysis 2.0 User Manual. P-value adjustment is performed using the Benjamini-Hochberg method of estimating false discovery rates (FDR). Clustering of genes for the final heatmap of differentially expressed genes was done using the Partitioning Around Medoids method using the fpc R library(Hennig, n.d.) that takes into consideration the direction and type of all signals on a pathway, and the position, role and type of every gene. Hypergeometric distribution was used to analyze the enrichment of pathways, gene ontology, domain structure, and other ontologies. The topGO R library(Alexa and Rahnenfuhrer, n.d.) was used to determine local similarities and dependencies between GO terms in order to perform Elim pruning correction. Several database sources were referenced for enrichment analysis, including Interpro(Mitchell et al., 2019), NCBI(Geer et al., 2010), MsigDB(Liberzon et al., 2011; Subramanian et al., 2005), REACTOME (Fabregat et al., 2018), WikiPathways(Slenter et al., 2018). Enrichment was calculated relative to a set of background genes relevant for the experiment. Gene expression data was exported and heat maps were generated in GraphPad prism 9.

### RNA-sequencing analysis of SC dataset

An RNA-sequencing dataset (GSE121589) from NCBI’s Gene Expression Omnibus database was analyzed(Shcherbina et al., 2020). This data set includes bulk RNA-seq of isolated SCs from young (2-3 mo. old) and aged (22-24 mo. old) muscle from uninjured and from regenerating muscle at 1, 3, 5, and 7 dpi. The raw data files were reprocessed by trimming adapters and low quality reads using TrimGalore! followed by alignment to the Ensembl GRCm38 mouse genome using HiSat2(Kim et al., 2015). StringTie was used to determine the transcripts per million (TPM) of genes in each sample and DESeq2 was used to identify differentially expressed genes (DEGs)(Love et al., 2014; Pertea et al., 2016). Expression of DE genes identified with the nCounter panel were assessed. Highly regulated genes between quiescent SCs (day 0) and early-activated SCs (day 1 and 3 post injury) were indicated by significant (<0.05) change in TPM.

### Statistics

Statistical analysis was performed with GraphPad Prism software using appropriate tests and minimum of 95% confidence interval for significance. Specific details of the statistical tests are given in the figure legends.

## Author contributions

LAM designed, performed, and interpreted all experiments, wrote manuscript; DJ created the MuSK^ΔIg3/ΔIg3^ (ΔIg3-MuSK) and *MuSK-Ig3^loxP/loxP^* mice; IC analyzed data, edited the manuscript and created Fig. 7. JRF designed and interpreted experiments, wrote manuscript.

## Acknowledgements

We thank Beth McKechnie for expert technical assistance and Youngwook Ahn of the Brown Transgenic core for generating the ΔIg3-MuSK mice. We thank K. O’Connor for generously providing the anti-MuSK monoclonal antibody and M. Huibers for the MuSK-MG patient IgG4 fraction. We thank G. Williams and the Brown Leduc Bioimaging Facility, K. Carlson & the Brown Flow Cytometry and Cell Sorter Facility, and C. Schorl and the Brown Genomics Facility for experimental assistance. Support: Brown University COBRE Center for Computational Biology: P20 GM10903; Mouse Transgenic and Gene Targeting Facility: P30 GM103410; Genomic Facility: P30 RR031153, P20 RR018728, S10 RR02763, NSF EPSCoR 0554548; DJ: 4R25GM083270 and 2T32AG041688; JRF: NIH awards U01 NS064295, R41 AG073144, R21 NS112743, R21 AG073743, and ALS Finding a Cure.

## Data Availability

The primary data, mouse models and reagents related to this work will be available to interested researchers upon reasonable request.

## Supplemental material

**Supplemental Figure 1.**
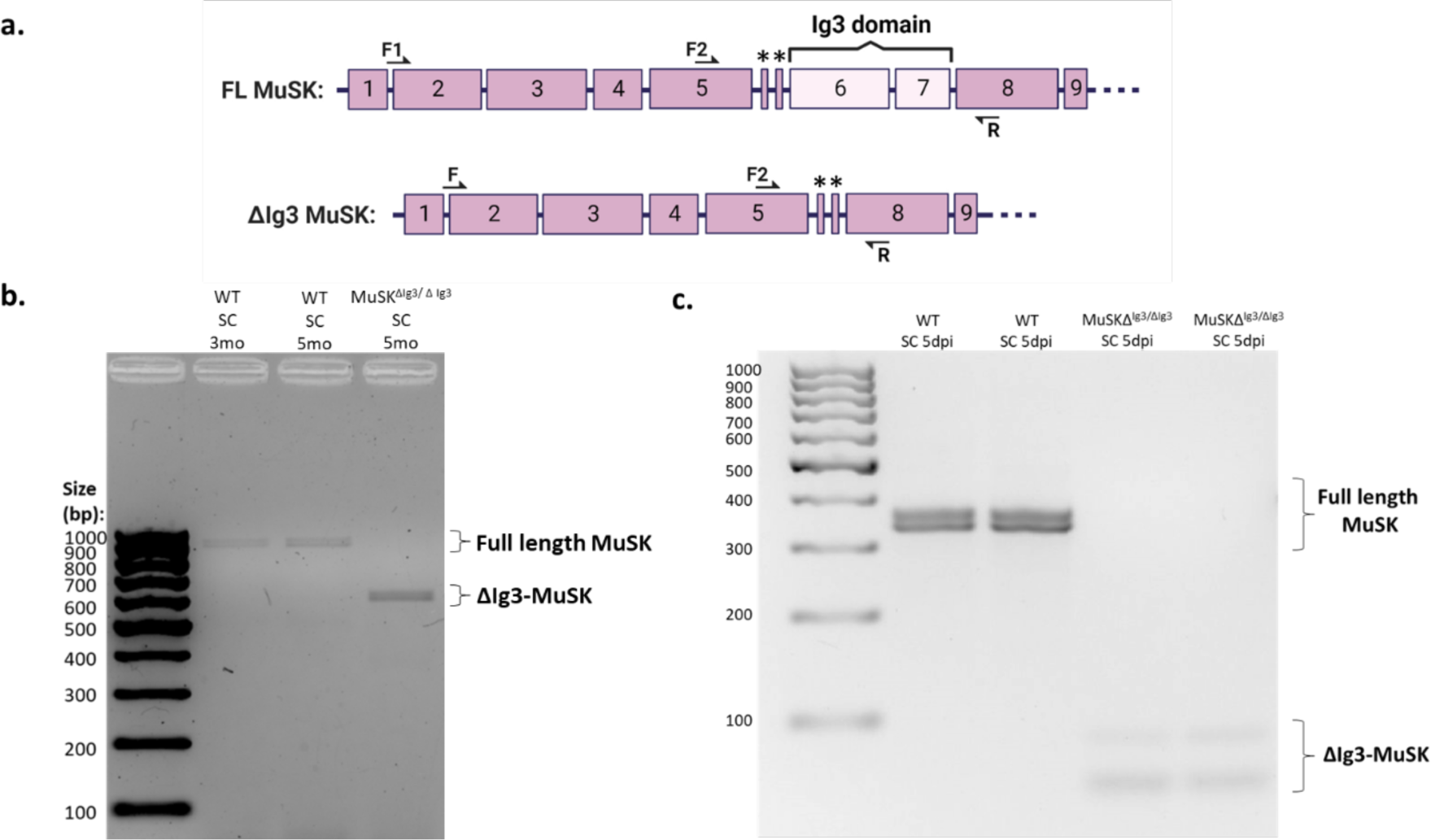
MuSK expression in SCs from uninjured and regenerating WT and ΔIg3-MuSK muscle. (a) Exon organization of MuSK. The two alternatively-spliced 30nt exons (5a and 5b) are indicated by asterisks. The PCR primers used for panels (b) and (c) are denoted F1 and F2, respectively. (**b.**) SCs from 3- and 5- month-old uninjured WT muscle express FL-MuSK; ΔIg3-MuSK is not detected. SCs from resting muscle from constitutive ΔIg3-MuSK mice express only ΔIg3-MuSK, as predicted. (**c.**) SCs from regenerating 5-month-old WT muscle 5 dpi express FL-MUSK, but no detectable ΔIg3-MuSK. SCs from regenerating muscle in ΔIg3-MuSK mice 5 dpi express ΔIg3-MuSK transcripts. Note that MuSK has two alternatively-spliced 30 nucleotide exons (5a and 5b) between exons 5 and 6 (Fig. 1a). Alternative splicing of the 5b exon is the basis for the multiple bands detected(Jaime et al., 2024).

**Supplemental Figure 2.**
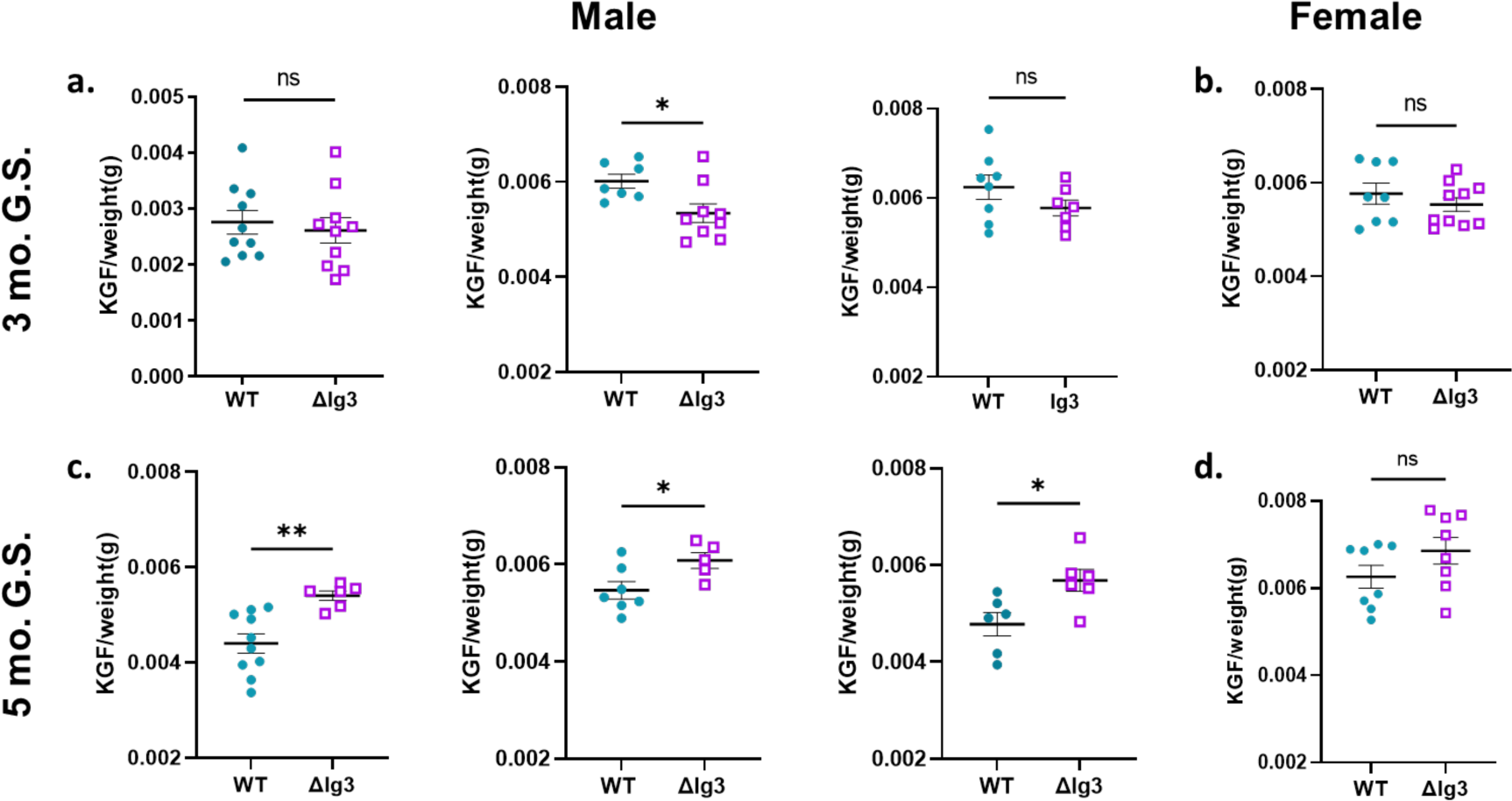
Grip strength in ΔIg3-MuSK and WT mice at 3 and 5 months of age in multiple cohorts. (**a**) Three different cohorts of male mice were used to assess 3 mo. grip strength (t-test, p=0.64, p=0.02, p=0.18). (**b**) One cohort of female mice was used to assess 3 mo. grip strength (t-test, p=0.38) (**c)** Three separate cohorts of male mice were used to assess 5 mo. grip strength (t-test, p=0.029,p=0.036, p=0.021). (**d**) One cohort of female mice was used to assess 5 mo. grip strength (t-test, p=0.16). (Note that left panels in a. and c. are the same data presented in the Fig. 2d and are shown here for comparison.)

**Supplemental Figure 3.**
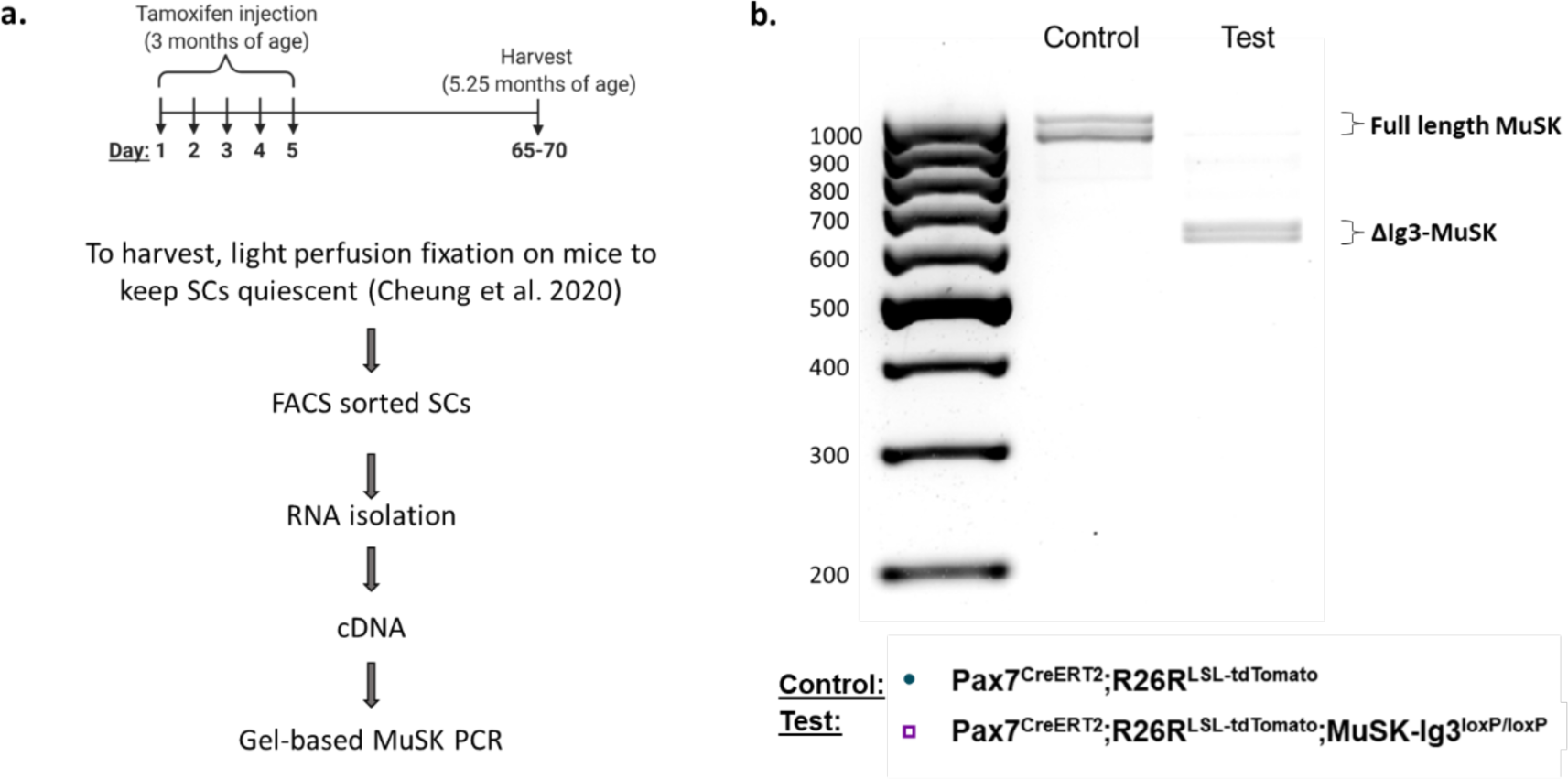
Tamoxifen treatment at 3 months induces Cre-mediated recombination of the MuSK-Ig3 domain in SCs from *Pax7^CreERT2^;R26R^LSL-tdTomato^;MuSK-Ig3^loxP/loxP^*mice. (**a.**) Experimental design. *Pax7^CreERT2^;R26R^LSL-tdTomato^;MuSK-Ig3^loxP/loxP^* and *Pax7^CreERT2^;R26R^LSL-tdTomato^* mice were injected with 5 daily doses of tamoxifen at 3 months of age and the muscles harvested 2 months later. Prior to muscle harvest mice were lightly perfusion fixed with 0.5% PFA to capture SCs in their *in vivo* state. RNA was isolated from FACS-purified SCs. (**b.**) Gel-based PCR analysis of FL-MuSK and ΔIg3-MuSK expression using the F1 and R primers (see Supp. Fig 1a.). Only FL-MuSK is detected in SCs from *Pax7^CreERT2^;R26R^LSL-tdTomato^* mice and only ΔIg3-MuSK (617 and 587bp) is detected in control *Pax7^CreERT2^;R26R^LSL-tdTomato^;MuSK-Ig3^loxP/loxP^* mice. See also Supp. Fig. 1 for details on primer location and band identification.

**Supplemental Fig. 4.**
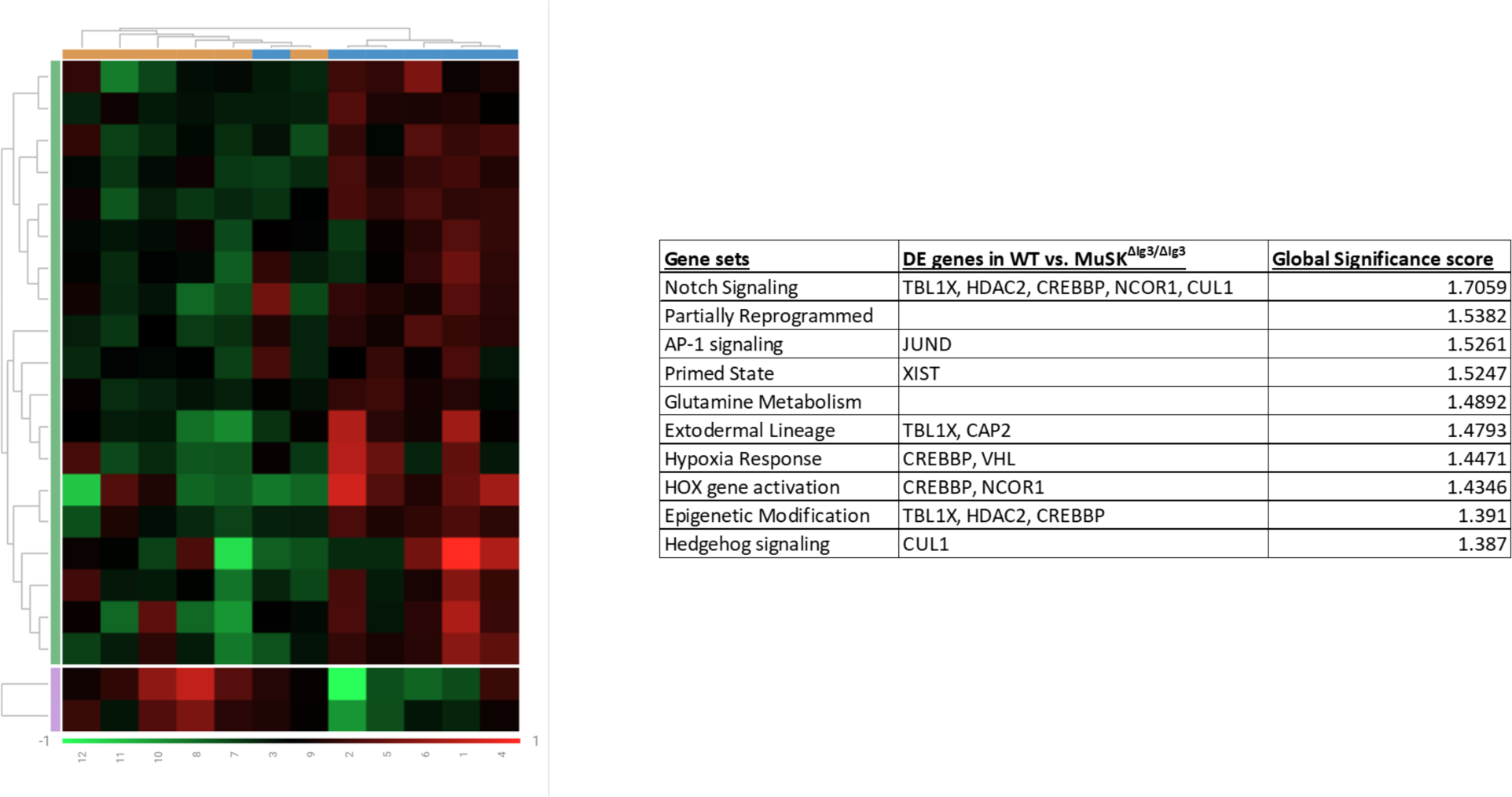
Nanostring nCounter unsupervised results. Heatmap showing the unsupervised hierarchical clustering of WT SCs (orange, n=6) and ΔIg3-MuSK SCs (blue, n=6) generated via analysis of NanoString Stem Cell Characterization panel on Rosalind. (**b)** Ranking of gene set functions according to the Global Significance Scores quantified by Nanostring software, based on unsupervised clustering.

**Supplemental Figure 5.**
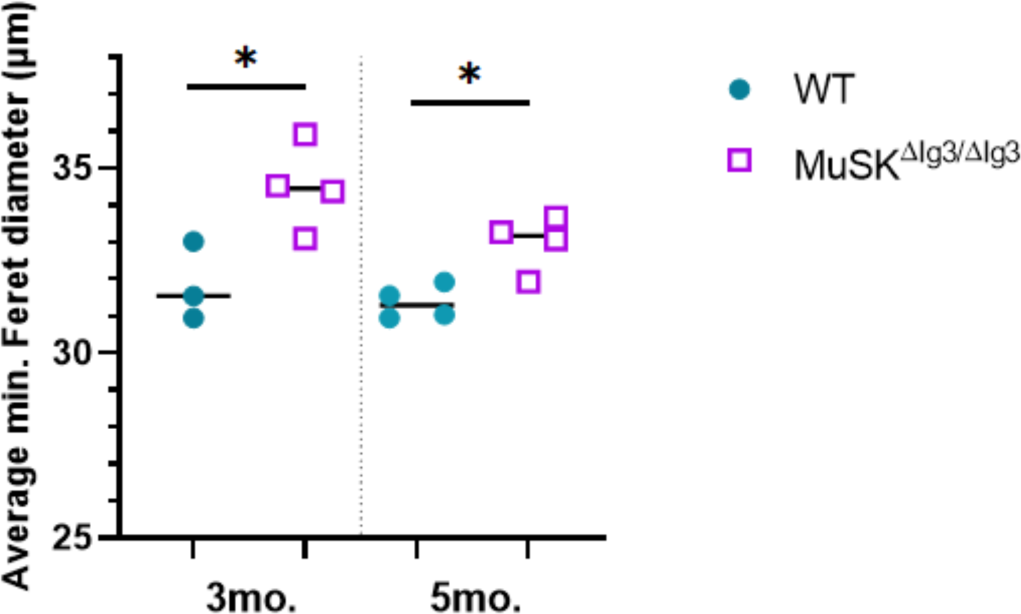
Female ΔIg3-MuSK TA myofibers are larger than WT at 3 and at 5 months. At 3 months and 5 months, the minimum Feret diameter is about 10% larger in ΔIg3-MuSK mouse muscle compared to WT (n=3-4 female mice, unpaired t-test, 3 mo. p=0.03; 5 mo. p=0.01)

## References

Ancel, S., Stuelsatz, P., Feige, J.N., 2021. Muscle Stem Cell Quiescence: Controlling Stemness by Staying Asleep. Trends Cell Biol. 10.1016/j.tcb.2021.02.006

Bachman, J.F., Chakkalakal, J.V., 2022. Insights into muscle stem cell dynamics during postnatal development. FEBS J. 289, 2710–2722.

Bachman, J.F., Klose, A., Liu, W., Paris, N.D., Blanc, R.S., Schmalz, M., Knapp, E., Chakkalakal, J.V., 2018. Prepubertal skeletal muscle growth requires Pax7-expressing satellite cell-derived myonuclear contribution. Development 145. 10.1242/dev.167197

Baghdadi, M.B., Castel, D., Machado, L., Fukada, S.-I., Birk, D.E., Relaix, F., Tajbakhsh, S., Mourikis, P., 2018. Reciprocal signalling by Notch-Collagen V-CALCR retains muscle stem cells in their niche. Nature 557, 714–718.

Bikou, A., Dermiki-Gkana, F., Penteris, M., Constantinides, T.K., Kontogiorgis, C., 2024. A systematic review of the effect of semaglutide on lean mass: insights from clinical trials. Expert Opin. Pharmacother. 25, 611–619.

Bjornson, C.R.R., Cheung, T.H., Liu, L., Tripathi, P.V., Steeper, K.M., Rando, T.A., 2012. Notch signaling is necessary to maintain quiescence in adult muscle stem cells. Stem Cells 30, 232–242.

Blau, H.M., Cosgrove, B.D., Ho, A.T.V., 2015. The central role of muscle stem cells in regenerative failure with aging. Nat. Med. 21, 854–862.

Bowen, D.C., Park, J.S., Bodine, S., Stark, J.L., Valenzuela, D.M., Stitt, T.N., Yancopoulos, G.D., Lindsay, R.M., Glass, D.J., DiStefano, P.S., 1998. Localization and regulation of MuSK at the neuromuscular junction. Dev. Biol. 199, 309–319.

Brack, A.S., Bildsoe, H., Hughes, S.M., 2005. Evidence that satellite cell decrement contributes to preferential decline in nuclear number from large fibres during murine age-related muscle atrophy. J. Cell Sci. 118, 4813–4821.

Briguet, A., Courdier-Fruh, I., Foster, M., Meier, T., Magyar, J.P., 2004. Histological parameters for the quantitative assessment of muscular dystrophy in the mdx-mouse. Neuromuscul. Disord. 14, 675–682.

Brunet, A., Goodell, M.A., Rando, T.A., 2023. Ageing and rejuvenation of tissue stem cells and their niches. Nat. Rev. Mol. Cell Biol. 24, 45–62.

Chamberlain, J.S., Olwin, B.B., 2010. Prevention of muscle aging by myofiber-associated satellite cell transplantation. Sci. Transl. Med.

Cosgrove, B.D., Gilbert, P.M., Porpiglia, E., Mourkioti, F., Lee, S.P., Corbel, S.Y., Llewellyn, M.E., Delp, S.L., Blau, H.M., 2014. Rejuvenation of the muscle stem cell population restores strength to injured aged muscles. Nat. Med. 20, 255–264.

Cramer, A.A.W., Prasad, V., Eftestøl, E., Song, T., Hansson, K.-A., Dugdale, H.F., Sadayappan, S., Ochala, J., Gundersen, K., Millay, D.P., 2020. Nuclear numbers in syncytial muscle fibers promote size but limit the development of larger myonuclear domains. Nat. Commun. 11, 6287.

de Morree, A., Rando, T.A., 2023. Regulation of adult stem cell quiescence and its functions in the maintenance of tissue integrity. Nat. Rev. Mol. Cell Biol. 10.1038/s41580-022-00568-6

Díaz-Moreno, M., Armenteros, T., Gradari, S., Hortigüela, R., García-Corzo, L., Fontán-Lozano, Á., Trejo, J.L., Mira, H., 2018. Noggin rescues age-related stem cell loss in the brain of senescent mice with neurodegenerative pathology. Proc. Natl. Acad. Sci. U. S. A. 115, 11625–11630.

Egner, I.M., Bruusgaard, J.C., Gundersen, K., 2016. Satellite cell depletion prevents fiber hypertrophy in skeletal muscle. Development 143, 2898–2906.

Eliazer, S., Muncie, J.M., Christensen, J., Sun, X., D’Urso, R.S., Weaver, V.M., Brack, A.S., 2019. Wnt4 from the Niche Controls the Mechano-Properties and Quiescent State of Muscle Stem Cells. Cell Stem Cell 25, 654–665.e4.

Fabregat, A., Jupe, S., Matthews, L., Sidiropoulos, K., Gillespie, M., Garapati, P., Haw, R., Jassal, B., Korninger, F., May, B., Milacic, M., Roca, C.D., Rothfels, K., Sevilla, C., Shamovsky, V., Shorser, S., Varusai, T., Viteri, G., Weiser, J., Wu, G., Stein, L., Hermjakob, H., D’Eustachio, P., 2018. The Reactome Pathway Knowledgebase. Nucleic Acids Res. 46, D649–D655.

Fish, L.A., Ewing, M.D., Jaime, D., Rich, K.A., Xi, C., Wang, X., Feder, R.E., Wharton, K.A., Rich, M.M., Arnold, W.D., Fallon, J.R., 2023. The MuSK-BMP pathway regulates synaptic Nav1.4 localization and muscle excitability. bioRxiv 10.1101/2023.05.17.541238 (J. Neuroscience, In Revision). 10.1101/2023.10.24.563837

Fish, L.A., Fallon, J.R., 2020. Multiple MuSK signaling pathways and the aging neuromuscular junction. Neurosci. Lett. 731, 135014.

Fuchs, E., Blau, H.M., 2020. Tissue Stem Cells: Architects of Their Niches. Cell Stem Cell 27, 532– 556.

Garcia-Osta, A., Tsokas, P., Pollonini, G., Landau, E.M., Blitzer, R., Alberini, C.M., 2006. MuSK expressed in the brain mediates cholinergic responses, synaptic plasticity, and memory formation. J. Neurosci. 26, 7919–7932.

García-Prat, L., Perdiguero, E., Alonso-Martín, S., Dell’Orso, S., Ravichandran, S., Brooks, S.R., Juan, A.H., Campanario, S., Jiang, K., Hong, X., Ortet, L., Ruiz-Bonilla, V., Flández, M., Moiseeva, V., Rebollo, E., Jardí, M., Sun, H.-W., Musarò, A., Sandri, M., Sol, A.D., Sartorelli, V., Muñoz-Cánoves, P., 2020. FoxO maintains a genuine muscle stem-cell quiescent state until geriatric age. Nat. Cell Biol. 10.1038/s41556-020-00593-7

Gattazzo, F., Laurent, B., Relaix, F., Rouard, H., Didier, N., 2020. Distinct Phases of Postnatal Skeletal Muscle Growth Govern the Progressive Establishment of Muscle Stem Cell Quiescence. Stem Cell Reports 15, 597–611.

Geer, L.Y., Marchler-Bauer, A., Geer, R.C., Han, L., He, J., He, S., Liu, C., Shi, W., Bryant, S.H., 2010. The NCBI BioSystems database. Nucleic Acids Res. 38, D492–6.

Gioftsidi, S., Relaix, F., Mourikis, P., 2022. The Notch signaling network in muscle stem cells during development, homeostasis, and disease. Skelet. Muscle 12, 9.

Girardi, F., Taleb, A., Ebrahimi, M., Datye, A., Gamage, D.G., Peccate, C., Giordani, L., Millay, D.P., Gilbert, P.M., Cadot, B., Le Grand, F., 2021. TGFβ signaling curbs cell fusion and muscle regeneration. Nat. Commun. 12, 750.

Goel, A.J., Rieder, M.-K., Arnold, H.-H., Radice, G.L., Krauss, R.S., 2017. Niche Cadherins Control the Quiescence-to-Activation Transition in Muscle Stem Cells. Cell Rep. 21, 2236–2250.

Goh, Q., Millay, D.P., 2017. Requirement of myomaker-mediated stem cell fusion for skeletal muscle hypertrophy. Elife 6. 10.7554/eLife.20007

Goh, Q., Song, T., Petrany, M.J., Cramer, A.A., Sun, C., Sadayappan, S., Lee, S.-J., Millay, D.P., 2019. Myonuclear accretion is a determinant of exercise-induced remodeling in skeletal muscle. Elife 8. 10.7554/eLife.44876

Gozo, M.C., Aspuria, P.-J., Cheon, D.-J., Walts, A.E., Berel, D., Miura, N., Karlan, B.Y., Orsulic, S., 2013. Foxc2 induces Wnt4 and Bmp4 expression during muscle regeneration and osteogenesis. Cell Death Differ. 20, 1031–1042.

Ham, D.J., Börsch, A., Chojnowska, K., Lin, S., Leuchtmann, A.B., Ham, A.S., Thürkauf, M., Delezie, J., Furrer, R., Burri, D., Sinnreich, M., Handschin, C., Tintignac, L.A., Zavolan, M., Mittal, N., Rüegg, M.A., 2022. Distinct and additive effects of calorie restriction and rapamycin in aging skeletal muscle. Nat. Commun. 13, 2025.

Hesser, B.A., Sander, A., Witzemann, V., 1999. Identification and characterization of a novel splice variant of MuSK. FEBS Lett. 442, 133–137.

Hogan, B.L., 1996. Bone morphogenetic proteins: multifunctional regulators of vertebrate development. Genes Dev. 10, 1580–1594.

Huijbers, M.G., Zhang, W., Klooster, R., Niks, E.H., Friese, M.B., Straasheijm, K.R., Thijssen, P.E., Vrolijk, H., Plomp, J.J., Vogels, P., Losen, M., Van der Maarel, S.M., Burden, S.J., Verschuuren, J.J., 2013. MuSK IgG4 autoantibodies cause myasthenia gravis by inhibiting binding between MuSK and Lrp4. Proc. Natl. Acad. Sci. U. S. A. 110, 20783–20788.

Jaime, D., Fish, L.A., Madigan, L.A., Xi, C., Piccoli, G., Ewing, M.D., Blaauw, B., Fallon, J.R., 2024. The MuSK-BMP pathway maintains myofiber size in slow muscle through regulation of Akt-mTOR signaling. Skelet. Muscle 14, 1.

Kann, A.P., Hung, M., Krauss, R.S., 2021. Cell–cell contact and signaling in the muscle stem cell niche. Curr. Opin. Cell Biol. 73, 78–83.

Keefe, A.C., Lawson, J.A., Flygare, S.D., Fox, Z.D., Colasanto, M.P., Mathew, S.J., Yandell, M., Kardon, G., 2015. Muscle stem cells contribute to myofibres in sedentary adult mice. Nat. Commun. 6, 7087.

Kim, D., Langmead, B., Salzberg, S.L., 2015. HISAT: a fast spliced aligner with low memory requirements. Nat. Methods 12, 357–360.

Kimmel, J.C., Hwang, A.B., Scaramozza, A., Marshall, W.F., Brack, A.S., 2020. Aging induces aberrant state transition kinetics in murine muscle stem cells. Development 147. 10.1242/dev.183855

Lee, S.-J., 2022. Myostatin: A Skeletal Muscle Chalone. Annu. Rev. Physiol. 10.1146/annurev-physiol-012422-112116

Lehallier, B., Gate, D., Schaum, N., Nanasi, T., Lee, S.E., Yousef, H., Moran Losada, P., Berdnik, D., Keller, A., Verghese, J., Sathyan, S., Franceschi, C., Milman, S., Barzilai, N., Wyss-Coray, T., 2019. Undulating changes in human plasma proteome profiles across the lifespan. Nat. Med. 25, 1843–1850.

Liberzon, A., Subramanian, A., Pinchback, R., Thorvaldsdóttir, H., Tamayo, P., Mesirov, J.P., 2011. Molecular signatures database (MSigDB) 3.0. Bioinformatics 27, 1739–1740.

Liu, W., Klose, A., Forman, S., Paris, N.D., Wei-LaPierre, L., Cortés-Lopéz, M., Tan, A., Flaherty, M., Miura, P., Dirksen, R.T., Chakkalakal, J.V., 2017. Loss of adult skeletal muscle stem cells drives age-related neuromuscular junction degeneration. Elife 6. 10.7554/eLife.26464

Love, M.I., Huber, W., Anders, S., 2014. Moderated estimation of fold change and dispersion for RNA-seq data with DESeq2. Genome Biol. 15, 550.

Massagué, J., 2012. TGFβ signalling in context. Nat. Rev. Mol. Cell Biol. 13, 616–630.

Mira, H., Andreu, Z., Suh, H., Lie, D.C., Jessberger, S., Consiglio, A., San Emeterio, J., Hortigüela, R., Marqués-Torrejón, M.A., Nakashima, K., Colak, D., Götz, M., Fariñas, I., Gage, F.H., 2010. Signaling through BMPR-IA regulates quiescence and long-term activity of neural stem cells in the adult hippocampus. Cell Stem Cell 7, 78–89.

Mitchell, A.L., Attwood, T.K., Babbitt, P.C., Blum, M., Bork, P., Bridge, A., Brown, S.D., Chang, H.-Y., El-Gebali, S., Fraser, M.I., Gough, J., Haft, D.R., Huang, H., Letunic, I., Lopez, R., Luciani, A., Madeira, F., Marchler-Bauer, A., Mi, H., Natale, D.A., Necci, M., Nuka, G., Orengo, C., Pandurangan, A.P., Paysan-Lafosse, T., Pesseat, S., Potter, S.C., Qureshi, M.A., Rawlings, N.D., Redaschi, N., Richardson, L.J., Rivoire, C., Salazar, G.A., Sangrador-Vegas, A., Sigrist, C.J.A., Sillitoe, I., Sutton, G.G., Thanki, N., Thomas, P.D., Tosatto, S.C.E., Yong, S.-Y., Finn, R.D., 2019. InterPro in 2019: improving coverage, classification and access to protein sequence annotations. Nucleic Acids Res. 47, D351–D360.

Mori, Shuuichi, Suzuki, S., Konishi, T., Kawaguchi, N., Kishi, M., Kuwabara, S., Ishizuchi, K., Zhou, H., Shibasaki, F., Tsumoto, H., Omura, T., Miura, Y., Mori, Seijiro, Higashihara, M., Murayama, S., Shigemoto, K., 2023. Proteolytic ectodomain shedding of muscle-specific tyrosine kinase in myasthenia gravis. Exp. Neurol. 361, 114300.

Murach, K.A., Fry, C.S., Dupont-Versteegden, E.E., McCarthy, J.J., Peterson, C.A., 2021. Fusion and beyond: Satellite cell contributions to loading-induced skeletal muscle adaptation. FASEB J. 35, e21893.

Murphy, K.T., Ryall, J.G., Snell, S.M., Nair, L., Koopman, R., Krasney, P.A., Ibebunjo, C., Holden, K.S., Loria, P.M., Salatto, C.T., Lynch, G.S., 2010. Antibody-directed myostatin inhibition improves diaphragm pathology in young but not adult dystrophic mdx mice. Am. J. Pathol. 176, 2425–2434.

Murphy, M.M., Lawson, J.A., Mathew, S.J., Hutcheson, D.A., Kardon, G., 2011. Satellite cells, connective tissue fibroblasts and their interactions are crucial for muscle regeneration. Development 138, 3625–3637.

Ono, Y., Calhabeu, F., Morgan, J.E., Katagiri, T., Amthor, H., Zammit, P.S., 2011. BMP signalling permits population expansion by preventing premature myogenic differentiation in muscle satellite cells. Cell Death Differ. 18, 222–234.

Pawlikowski, B., Pulliam, C., Betta, N.D., Kardon, G., Olwin, B.B., 2015. Pervasive satellite cell contribution to uninjured adult muscle fibers. Skelet. Muscle 5, 42.

Pawlikowski, B., Vogler, T.O., Gadek, K., Olwin, B.B., 2017. Regulation of skeletal muscle stem cells by fibroblast growth factors. Dev. Dyn. 246, 359–367.

Perkins, J.R., Dawes, J.M., McMahon, S.B., Bennett, D.L.H., Orengo, C., Kohl, M., 2012. ReadqPCR and NormqPCR: R packages for the reading, quality checking and normalisation of RT-qPCR quantification cycle (Cq) data. BMC Genomics 13, 296.

Pertea, M., Kim, D., Pertea, G.M., Leek, J.T., Salzberg, S.L., 2016. Transcript-level expression analysis of RNA-seq experiments with HISAT, StringTie and Ballgown. Nat. Protoc. 11, 1650–1667.

Petrany, M.J., Swoboda, C.O., Sun, C., Chetal, K., Chen, X., Weirauch, M.T., Salomonis, N., Millay, D.P., 2020. Single-nucleus RNA-seq identifies transcriptional heterogeneity in multinucleated skeletal myofibers. Nat. Commun. 11, 6374.

Rodgers, J.T., King, K.Y., Brett, J.O., Cromie, M.J., Charville, G.W., Maguire, K.K., Brunson, C., Mastey, N., Liu, L., Tsai, C.-R., Goodell, M.A., Rando, T.A., 2014. mTORC1 controls the adaptive transition of quiescent stem cells from G0 to G(Alert). Nature 510, 393–396.

Sambasivan, R., Yao, R., Kissenpfennig, A., Van Wittenberghe, L., Paldi, A., Gayraud-Morel, B., Guenou, H., Malissen, B., Tajbakhsh, S., Galy, A., 2011. Pax7-expressing satellite cells are indispensable for adult skeletal muscle regeneration. Development 138, 3647–3656.

Shavlakadze, T., Morris, M., Fang, J., Wang, S.X., Zhu, J., Zhou, W., Tse, H.W., Mondragon-Gonzalez, R., Roma, G., Glass, D.J., 2019. Age-Related Gene Expression Signature in Rats Demonstrate Early, Late, and Linear Transcriptional Changes from Multiple Tissues. Cell Rep. 28, 3263–3273.e3.

Shcherbina, A., Larouche, J., Fraczek, P., Yang, B.A., Brown, L.A., Markworth, J.F., Chung, C.H., Khaliq, M., de Silva, K., Choi, J.J., Fallahi-Sichani, M., Chandrasekaran, S., Jang, Y.C., Brooks, S.V., Aguilar, C.A., 2020. Dissecting Murine Muscle Stem Cell Aging through Regeneration Using Integrative Genomic Analysis. Cell Rep. 32, 107964.

Siegel, A.L., Kuhlmann, P.K., Cornelison, D.D.W., 2011.Muscle satellite cell proliferation and association: new insights from myofiber time-lapse imaging. Skelet. Muscle 1, 7.

Slenter, D.N., Kutmon, M., Hanspers, K., Riutta, A., Windsor, J., Nunes, N., Mélius, J., Cirillo, E., Coort, S.L., Digles, D., Ehrhart, F., Giesbertz, P., Kalafati, M., Martens, M., Miller, R., Nishida, K., Rieswijk, L., Waagmeester, A., Eijssen, L.M.T., Evelo, C.T., Pico, A.R., Willighagen, E.L., 2018. WikiPathways: a multifaceted pathway database bridging metabolomics to other omics research. Nucleic Acids Res. 46, D661–D667.

Sousa-Victor, P., García-Prat, L., Muñoz-Cánoves, P., 2022. Control of satellite cell function in muscle regeneration and its disruption in ageing. Nat. Rev. Mol. Cell Biol. 23, 204–226.

Stantzou, A., Schirwis, E., Swist, S., Alonso-Martin, S., Polydorou, I., Zarrouki, F., Mouisel, E., Beley, C., Julien, A., Le Grand, F., Garcia, L., Colnot, C., Birchmeier, C., Braun, T., Schuelke, M., Relaix, F., Amthor, H., 2017. BMP signaling regulates satellite cell-dependent postnatal muscle growth. Development 144, 2737–2747.

Stiegler, A.L., Burden, S.J., Hubbard, S.R., 2006. Crystal structure of the agrin-responsive immunoglobulin-like domains 1 and 2 of the receptor tyrosine kinase MuSK. J. Mol. Biol. 364, 424–433.

Subramanian, A., Tamayo, P., Mootha, V.K., Mukherjee, S., Ebert, B.L., Gillette, M.A., Paulovich, A., Pomeroy, S.L., Golub, T.R., Lander, E.S., Mesirov, J.P., 2005. Gene set enrichment analysis: a knowledge-based approach for interpreting genome-wide expression profiles. Proc. Natl. Acad. Sci. U. S. A. 102, 15545–15550.

Tabula Muris Consortium, Overall coordination, Logistical coordination, Organ collection and processing, Library preparation and sequencing, Computational data analysis, Cell type annotation, Writing group, Supplemental text writing group, Principal investigators, 2018. Single-cell transcriptomics of 20 mouse organs creates a Tabula Muris. Nature 562, 367– 372.

Tunc-Ozcan, E., Brooker, S.M., Bonds, J.A., Tsai, Y.-H., Rawat, R., McGuire, T.L., Peng, C.-Y., Kessler, J.A., 2021. Hippocampal BMP signaling as a common pathway for antidepressant action. Cell. Mol. Life Sci. 10.1007/s00018-021-04026-y

Wen, Y., Murach, K.A., Vechetti, I.J., Jr, Fry, C.S., Vickery, C., Peterson, C.A., McCarthy, J.J., Campbell, K.S., 2018. MyoVision: software for automated high-content analysis of skeletal muscle immunohistochemistry. J. Appl. Physiol. 124, 40–51.

White, R.B., Biérinx, A.-S., Gnocchi, V.F., Zammit, P.S., 2010. Dynamics of muscle fibre growth during postnatal mouse development. BMC Dev. Biol. 10, 21.

Yilmaz, A., Kattamuri, C., Ozdeslik, R.N., Schmiedel, C., Mentzer, S., Schorl, C., Oancea, E., Thompson, T.B., Fallon, J.R., 2016. MuSK is a BMP co-receptor that shapes BMP responses and calcium signaling in muscle cells. Sci. Signal. 9, ra87.

Yue, F., Bi, P., Wang, C., Shan, T., Nie, Y., Ratliff, T.L., Gavin, T.P., Kuang, S., 2017. Pten is necessary for the quiescence and maintenance of adult muscle stem cells. Nat. Commun. 8, 14328.

Yue, L., Cheung, T.H., 2020. Protocol for Isolation and Characterization of In Situ Fixed Quiescent Muscle Stem Cells. STAR Protoc 1, 100128.

Yue, L., Wan, R., Luan, S., Zeng, W., Cheung, T.H., 2020. Dek Modulates Global Intron Retention during Muscle Stem Cells Quiescence Exit. Dev. Cell 53, 661–676.e6.

Zammit, P.S., Heslop, L., Hudon, V., Rosenblatt, J.D., Tajbakhsh, S., Buckingham, M.E., Beauchamp, J.R., Partridge, T.A., 2002. Kinetics of myoblast proliferation show that resident satellite cells are competent to fully regenerate skeletal muscle fibers. Exp. Cell Res. 281, 39–49.

Zeng, W., Yue, L., Lam, K.S.W., Zhang, W., So, W.-K., Tse, E.H.Y., Cheung, T.H., 2022. CPEB1 directs muscle stem cell activation by reprogramming the translational landscape. Nat. Commun. 13, 947.

Zhang, L., Noguchi, Y.-T., Nakayama, H., Kaji, T., Tsujikawa, K., Ikemoto-Uezumi, M., Uezumi, A., Okada, Y., Doi, T., Watanabe, S., Braun, T., Fujio, Y., Fukada, S.-I., 2019. The CalcR-PKA-Yap1 Axis Is Critical for Maintaining Quiescence in Muscle Stem Cells. Cell Rep. 29, 2154–2163.e5.

Zhang, Y., Lahmann, I., Baum, K., Shimojo, H., Mourikis, P., Wolf, J., Kageyama, R., Birchmeier, C., 2021. Oscillations of Delta-like1 regulate the balance between differentiation and maintenance of muscle stem cells. Nat. Commun. 12, 1318.

Zong, Y., Zhang, B., Gu, S., Lee, K., Zhou, J., Yao, G., Figueiredo, D., Perry, K., Mei, L., Jin, R., 2012. Structural basis of agrin-LRP4-MuSK signaling. Genes Dev. 26, 247–258.

